# A synthetic oligo library and sequencing approach reveals an insulation mechanism encoded within bacterial σ^54^ promoters

**DOI:** 10.1101/086108

**Authors:** Lior Levy, Leon Anavy, Oz Solomon, Roni Cohen, Michal Brunwasser-Meirom, Shilo Ohayon, Orna Atar, Sarah Goldberg, Zohar Yakhini, Roee Amit

## Abstract

We use an oligonucleotide library of over 10000 variants together with a synthetic biology approach to identify an insulation mechanism encoded within a subset of σ^54^ promoters. Insulation manifests itself as dramatically reduced protein expression for a downstream gene that may be expressed by transcriptional read-through. The insulation we observe is strongly associated with the presence of short CT-rich motifs (3-5 bp), positioned within 25 bp upstream of the Shine-Dalgarno (SD) motif of the silenced gene. We hypothesize that insulation is effected by binding of the RBS to the upstream CT-rich motif. We provide evidence to support this hypothesis using mutations to the CT-rich motif and gene expression measurements on multiple sequence variants. Modelling is also consistent with this hypothesis. We show that the strength of the silencing, effected by insulation, depends on the location and number of CT-rich motifs encoded within the promoters. Finally, we show that in *E.coli* these insulator sequences are preferentially encoded within σ^54^ promoters as compared to other promoter types, suggesting a regulatory role for these sequences in natural contexts. Our findings suggest that context-related regulatory effects may often be due to sequence-specific interactions encoded sparsely by short motifs that are not easily detected by lower throughput studies. Such short sequence-specific phenomena can be uncovered with a focused OL design that filters out the sequence noise, as exemplified herein.

## Introduction

Deconstructing genomes to their basic parts and then using those parts to construct *de novo* gene regulatory architectures are central hallmarks of synthetic biology. As a first step, a thorough breakdown of a genome to its basic regulatory and functional elements is required. Then, each element can be analyzed to decipher the properties and mechanisms that drive and attenuate its activity. Lastly, well-defined and well-characterized elements can, in theory, be used as building blocks for *de novo* systems. However, in practice, *de novo* genetic systems often fail to operate as designed, due to the complex interplay between different supposedly well-characterized elements.

A possible cause of such unexpected behavior is context. Here, *context* refers to the DNA sequences that connect the different elements of the *de novo* circuit, the flanking segments within the elements, and even parts of particular elements, any of which may encode unknown regulatory roles. Often, context effects are due to short-range sequence-based interactions with nearby elements (Korbel et al., 2004). Such interactions might confer some secondary regulatory effect that is overlooked by standard analysis methods or is masked by a stronger regulatory effect in the native setting (Farley et al., 2015). Context effects can emerge from RNA secondary structure, or from larger scale genomic properties that involve nearby transcriptionally active loci. For instance, the formation of secondary structure either near the ribosome binding sites, or in configurations that sequester the RBS via hybridization by an anti-Shine Dalgarno (aSD) sequence have been suggested as strongly inhibiting or modulating the initiation of translation (Campo et al., 2015; De Smit and Duin, 1990; Ma et al., 2002; Schwartz et al., 1981). In bacteria, context effects have also been explored with respect to coding regions. For example, bacterial codon usage 30 nt downstream of the start codon has been shown to be biased towards unstable secondary structure and is generally GC-poor as a result (Bentele et al., 2013; Gu et al., 2010; Kudla et al., 2009). Other intragenic regulatory phenomena that have been recently proposed in bacteria involve inactive σ^54^ promoters (i.e. promoters that do not have an associated upstream activating sequence that binds enhancer proteins) that instead of triggering expression, function as binding sites for large DNA binding proteins that repress expression either internally within genes or by competing with the binding of transcriptionally active RNAP complexes (Bonocora et al., 2015; J. Schaefer et al., 2015). Alternatively, dynamical processes such as transcriptional interference by an incoming RNA polymerase, or transcriptional read-through from an upstream locus can also alter gene expression in a way that is not encoded in the individual parts (Epshtein et al., 2003; Hao et al., 2016; Shearwin et al., 2005). In summary, in light of such diverse effects, a more systematic understanding of context-related regulatory mechanisms, their effectors and their sequence related determinants is needed. Such understanding is important both in the context of natural cellular processes and for the design of reliable synthetic biology modalities.

Directed evolution screens have been suggested as a technique to avoid unwanted context effects in synthetic constructs (Yokobayashi et al., 2002). These do not fit every scenario and are often impractical for actual circuit designs. Synthetic oligonucleotide libraries (OLs), together with high-throughput focused screening methods, provide an alternative approach that enables direct investigation of context-related effects. Synthetic OLs have been used to examine regulatory elements systematically, and have revealed the effects of element location and multiplicity (Kinney et al., 2010; Shalem et al., 2015; Sharon et al., 2012, 2014). In this work, we applied OL technology to investigate secondary context-related phenomena in bacterial non-coding regulatory elements. An OL of closely related variants was designed and then embedded in a synthetic transcriptional read-through regulatory architecture to explore the underlying sequence determinants of a down-regulation effect. Following the experimental scheme introduced by (Sharon et al., 2012), the library was first sorted using flow cytometry into bins, which are subsequently sequenced. The combined sequencing and fluorescence data set facilitates the extraction of individual expression distributions for each variant. As a result, this process, which has been referred to as sort-seq or flow-seq, (see (Peterman and Levine, 2016) for review) generates a large distribution of regulatory elements and their associated behavior.

In this study, we used sort-seq to investigate a transcriptional insulation phenomenon, first observed in smaller scale within the context of the *glnK* σ^54^ promoter (glnKp). In the case of our genetic circuit design, RNA of a downstream gene cannot be transcribed by the σ^54^ promoter itself, but rather via transcriptional read-through. Namely, by an RNA-polymerase arriving from an upstream locus. To understand the differential silencing of a downstream mCherry protein we designed the sort-seq library to explore hundreds of mutant variants of the original *E. coli* glnKp context, and search for similar insulator contexts in other annotated σ^54^-promoters from multiple bacterial species. Using this focused OL we found that silenced variants are associated with short 3-5 nucleotide CT-rich segments within the σ^54^-promoters. Furthermore, our analysis revealed that the strength of silencing is associated with the number and the location of the short CT-motifs relative to the σ^54^-promoter transcriptional start site (TSS). Thus, our sort-seq analysis points to a bacterial insulation mechanism encoded not by a typical position specific scoring matrix (PSSM)-like motif, but rather through the presence of short sequence segments. CT-rich sequences are potentially complementary to the Shine-Dalganro (SD) sequence and we therefore hypothesize an anti-SD to SD binding as the mechanism that drives insulation and provide evidence to support this hypothesis.

## Results

### The σ^54^ glnK promoter silences expression from an upstream promoter

We engineered a set of synthetic circuits to test the components of bacterial enhancers in *E. coli*, initially in identical context. Bacterial enhancers typically consist of a poised σ^54^ (σ^A/C^ in gram-positive) promoter, an upstream activating sequence (UAS) consisting of a tandem of binding sites for some activator protein located 100-200 bp away (e.g. NtrC, PspF, LuxO, etc.), and an intervening sequence, facilitating DNA looping which often harbors additional transcription factor binding sites (Amit et al., 2011; Atkinson et al., 2002; Claverie-Martin and Magasanik, 1991; Kiupakis and Reitzer, 2002). In our study of enhancer components, each synthetic circuit consisted of a UAS element and a σ^54^ promoter that were taken out of their natural contexts and placed in an identical context, namely with the same 70 bp loop sequence between the UAS and the TSS of the promoter, and upstream of the same mCherry reporter gene (see Fig. 1A, Fig. S1, and Fig. S2). We chose five *E. coli* σ^54^ promoters of varying known strengths (Atkinson et al., 2002, 2003; Claverie-Martin and Magasanik, 1991; Feng et al., 1995; Kiupakis and Reitzer, 2002; Reitzer and Schneider, 2001) (glnAp2, glnKp, glnHp, astCp, nacp), and a no-promoter control. Ten UAS sequences were selected to cover a wide variety of binding affinities for NtrC and included four natural tandems, five chimeric tandems made from two halves of naturally occurring UASs, and one natural UAS, which is known to harbor a σ^70^ promoter overlapping the NtrC binding sites (glnAp1). Altogether, we synthesized 50 bacterial enhancers and 16 negative control circuits lacking either a UAS or a promoter. Finally, we compared mCherry levels between NtrC induction and non-induction states (see Methods for details of the positive-feedback synthetic enhancer circuit (Amit et al., 2011)).

**Figure 1:**
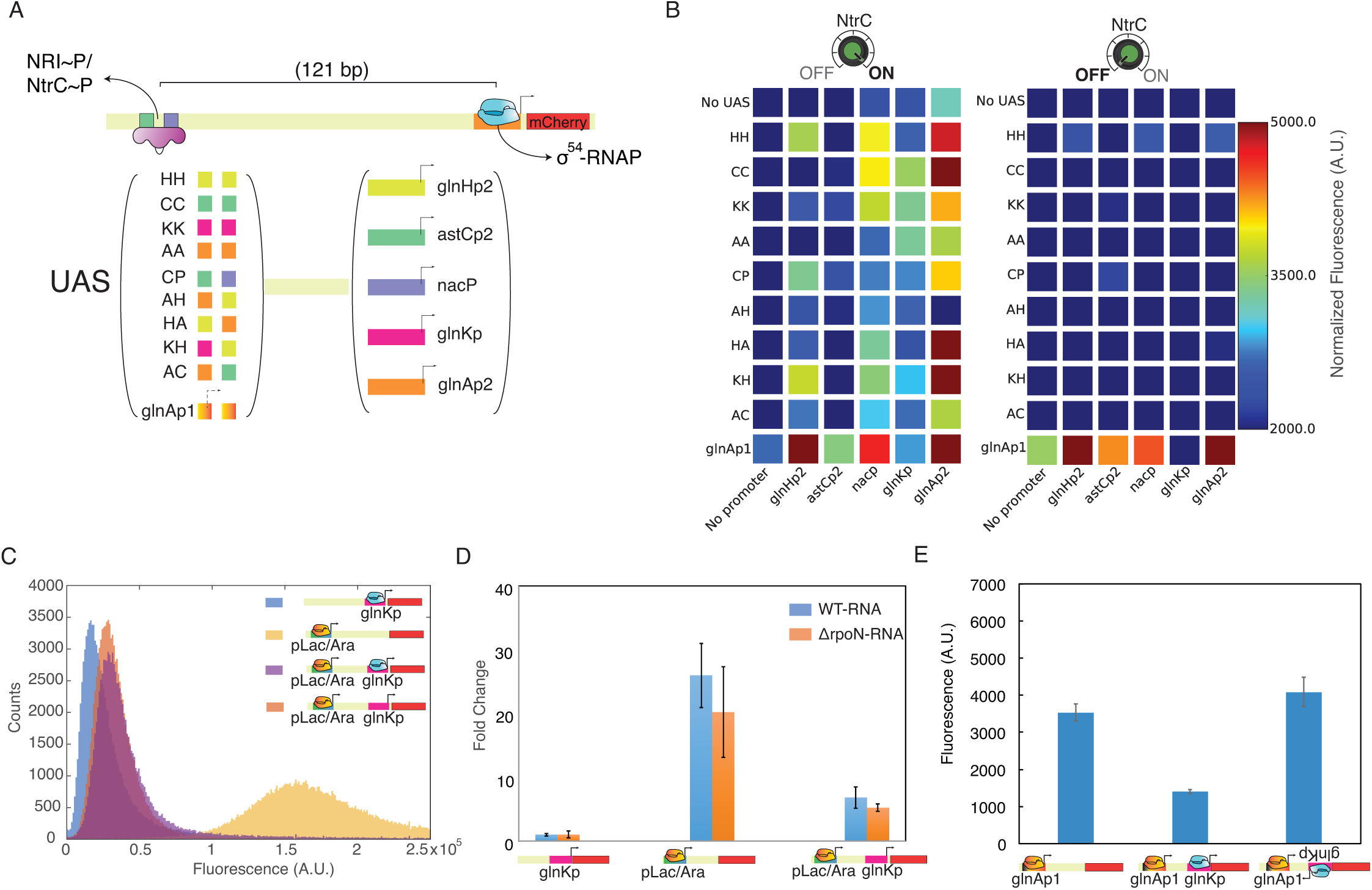
The glnKp σ^54^ promoter can down-regulate another promoter positioned upstream. (A) Synthetic enhancer design showing the different UAS and σ^54^ promoter combinations used in the experiment. (B) Left: mCherry expression with enhancer switched to “on’’ (NtrC induced), showing varying response for each promoter. Note that for the dual UAS-σ^70^ promoter glnAp1 there is expression with the “no promoter’’ control. Right: mCherry expression for enhancers switched to “off’’ (NtrC not induced), showing “on’’ behavior for all enhancers containing the dual UAS-σ^70^ promoter, except for the enhancer with the glnKp. Error level for the mean expression values are provided in Table S3. (C) Flow cytometry data comparing mCherry fluorescence for the glnKp strain in the *E. coli* TOP10 strain (purple) and in the σ^54^ knock-out strain (TOP10:Δ*rpoN*, orange). (D) qPCR data showing a reduction in mRNA level in the silenced strain (right) as compared with non-silenced strains (middle) and the no-σ^70^ control (left). (E) plate-reader data showing rescue of mCherry fluorescence when the orientation of the glnKp is flipped relative to the upstream σ^70^ promoter. Associated with Figure S1, Table S1, S2, and S3.

We report the mean fluorescence expression-level data in steady state together with their variation for the synthetic enhancers as Table S3. The left panel depicts mean mCherry expression levels with NtrC induced to high titers within the cells. The plot shows that all synthetic enhancer circuits are capable of generating fluorescence expression as compared with a no- σ^54^ -promoter control. The promoters which were previously reported to be "weak" (glnHp and astCp) and naturally bound by either IHF or ArgR (Claverie-Martin and Magasanik, 1991; Hoover et al., 1990; Kiupakis and Reitzer, 2002) were indeed found to generate lower levels of expression as compared with glnAp2, nacp, and glnKp (p-value<0.05, 10^−3^ paired t-test, for glnHp2 and astCp2, respectively). Variability of expression driven by glnAp2 is significantly higher than that of nacP and glnKp (p-value<0.01, F-test for variance equality). Finally, the glnAp1 UAS that contains an overlapping σ^70^ promoter induces expression in the no-promoter control, as expected.

To characterize the activity of the σ^70^ promoter in glnAp1 (the natural UAS for glnAp2), we plot the expression level data of the synthetic enhancers with NtrC uninduced in the right panel of Fig. 1B. Without NtrC, σ^54^ promoter expression should be silent and indeed, the only UAS for which we observed significant expression was glnAp1 (p-value<0.05, t-test after correcting for multiple testing). This is due to its dual role as a σ^70^ promoter in addition to being a σ^54^ UAS. However, glnAp1 showed a detectible fluorescence response for only four of the five promoters. The σ^54^ promoter glnKp manifested a different behavior. Namely, the glnAp1 UAS did not generate detectible expression with the glnK promoter, as compared with each of the other promoters (t-tests, using the distribution of technical replicates, p-value<0.01 for all promoters). Thus, there seems to be an inhibitory mechanism embedded within the σ^54^ promoter glnKp.

We initially reasoned that the inhibitory phenomenon might be explained by unusually tight binding of the σ^54^-RNAP complex to the glnKp core region, leading to the formation of a physical "road-block", which interferes with any upstream transcribing RNAP holoenzymes. To check this hypothesis, we constructed another gene circuit in which a pLac/Ara (σ^70^) promoter was placed upstream of the σ^54^ glnKp instead of the glnAp1 UAS. In Fig. 1C, we show that the circuit with both the pLac/Ara promoter and the glnKp (purple) generates about a factor of ten less fluorescence than the control lacking glnKp (yellow). However, when the circuit was placed in a Δ*rpoN* knockout strain (*rpoN* encodes the σ^54^ RNAP subunit), the same reduction in fluorescence was observed (orange). Moreover, in Fig. 1D (center and right bars) we show that the reduction was observed not only at the protein level, but also at the mRNA level, albeit to a lesser extent. The effect was observed only for glnKp oriented in the 5'-to-3' direction relative to pLac/Ara, as flipping the orientation of the 50 bp glnKp sequence abolished the inhibitory effect (Fig. 1E). Consequently, in the context of our construct, the glnKp sequence not only encodes a σ^54^ promoter, but also some function that leads to silencing and that is active when this sequence is placed downstream from an active σ^70^ promoter and upstream to the mCherry start codon. This silencing occurs both with and without rpoN.

### glnK promoter encodes an insulator sequence

In order to study a closer to nature configuration involving the glnK promoter, we encoded the full *glutathione S-transferase* (*GST*) gene upstream of the glnKp (under the control of pLac/Ara), with and without its own Shine-Dalgarno (SD) motif. By adding a gene upstream of the glnKp, we engineered a system that closely resembles a typical genomic architecture of one operon following another, thus allowing for transcriptional read-through of the downstream gene from the upstream promoter. We reasoned that a translated gene placed upstream of an aSD sequence would protect the entire mRNA from the pyrophosphotation of the 5’end by RppH (Deana et al., 2008). This, in turn, would inhibit the RnaseE degradation pathway (Mackie, 1998), leading to a partial rescue of the mCherry silencing effect. Previous studies (Calin-Jageman and Nicholson, 2003; Deana et al., 2008; Hui et al., 2014; Mackie, 1998; Richards et al., 2012; Robertson, 1982) have shown that a folded RNA state frequently triggers increased degradation of the untranslated RNA. Since we also observed reduced RNA levels in the silenced strain (Fig. 1D), we wanted to rule out that the possibility of the silencing effect being a degradation artifact of our original circuit design of two closely positioned promoters. The quantitative PCR (qPCR) results of the additional GST strains are shown in Fig. 2A. We plot data for four strains: glnKp variant without the *GST* gene (left bar), *GST*+glnKp+*mCherry* (second from the left), RBS+*GST*+glnKp+*mCherry* (second from the right), and a non-silenced strain (right). In Fig. 2A it can be seen that the mCherry mRNA level for the strain with a non-translated *GST* gene encoded upstream is identical to the one measured for the glnKp variant, and can thus be considered silenced. However, when the GST is translated, the mCherry mRNA levels rise considerably by a factor of ~3, representing approximately 50% recovery as compared with the non-silenced strain.

**Figure 2:**
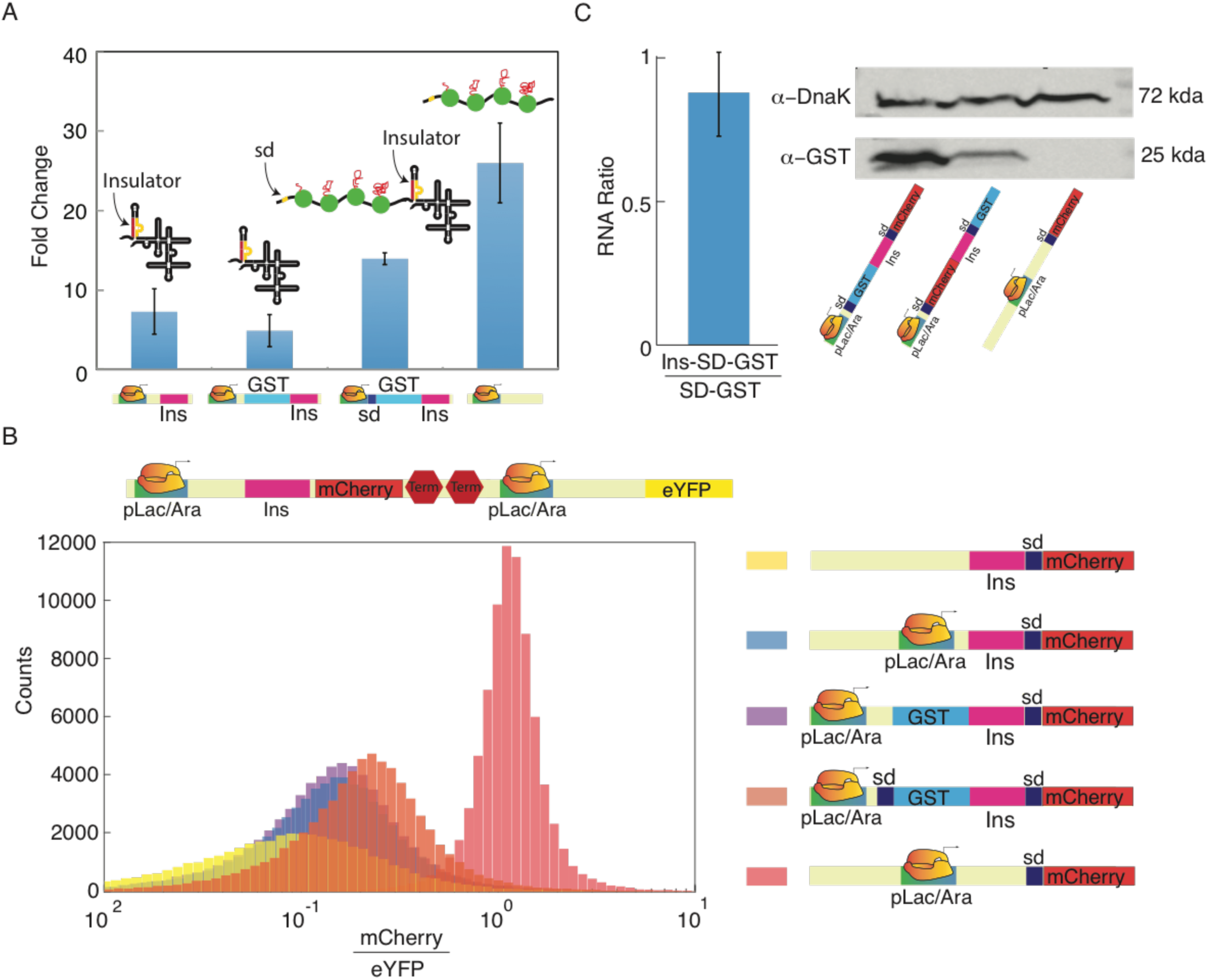
GlnKp down-regulation is generated by an aSD:SD interaction. (A) qPCR measurements showing the rescue of mCherry mRNA levels by a translated *GST* placed upstream of glnKp (third bar from left), as compared with a non-translated *GST* (second bar from left). The no-glnKp control is shown for comparison (fourth bar from left). (B) Flow cytometry data showing strong insulation despite adding the *gst* gene. (Top) Circuit diagram showing added eYFP module without an insulating component for mCherry to EYFP expression level ratio measurements (e.l.r). (Bottom) Data for the following constructs: no σ^70^ promoter control (yellow), glnKp construct (blue), glnKp construct with a *gst* gene encoded upstream with (orange) and without an RBS (purple) respectively, and the no- σ^54^ promoter control (red). (C) Insulation of a GST context. (Left) RT-PCR ratio of insulated and non-insulated GST RNA showing no effect. (Right) Western blot with α-GST comparing non-insulated (left band), insulated (middle band), and no-GST control (right band). Associated with Figure S2.

Next, we measured the recovery in expression level ratio of mCherry on the same strains. To get a quantitative assessment of the relative increase between the different strains, we added a non-insulated circuit expressing eYFP on the same plasmid (Fig. 2B-top), which allowed us to measure an mCherry to EYFP expression level ratio (e.l.r) that is less prone to expression level noise. In Fig. (2B-bottom), we plot the flow cytometry distributions for the following strains: the glnKp variant without an active upstream promoter (yellow), glnKp variant without *GST* (blue), *GST*+glnKp+*mCherry* (purple), RBS+*GST*+glnKp+mCherry (orange), and the variant with a non-silencing 5’UTR (red - note that in the legend the glnKp promoter is labelled in pink as “insulator”). A close examination of the data shows that an insignificant increase in e.l.r (Fig. 2C) is measured for the translationally active *GST* strain as compared with the e.l.r levels of the non-silenced strain. While this e.l.r level is consistent with the partial recovery in mRNA levels shown in Fig. 2A (two fold as compared with three fold), it remains smaller by about ten fold from the construct that does not encode the putative silencing 5’UTR sequence.

Given the recovery in mRNA levels and the lack thereof in e.l.r, we hypothesized that the silencing phenomenon we observed occurs at the post-transcriptional level. A close examination (Fig. S2) of the sequence encoded in the flanking region of glnKp reveals that there is a CU-rich segment located ~20 bp upstream of the SD sequence. This sequence can potentially form a hairpin structure with the sequence. RBS sequestration via hairpin formation has been previously implicated in gene silencing (Campo et al., 2015; De Smit and Duin, 1990; Ma et al., 2002; Schwartz et al., 1981). A hairpin presumably inhibits the formation of an elongating 70S ribosomal subunit in bacteria, thus leading to the silencing of the downstream gene. In our case, this aSD sequence is encoded within the glnK promoter and cannot be transcribed by the σ^54^ promoter itself. Instead, it can only become active and *insulate* the downstream σ^54^-regulated gene from translation, when the template mRNA is transcribed via transcriptional read-through based events. A visual representation of possible secondary structures (Hofacker et al., 1994) for the 5’UTR region of the constructs with the no σ^54^ promoter (top) and with the glnKp context (bottom) are shown in Fig. S2. The structure models suggest that while the RBS for the construct without σ^54^ promoter remains single stranded, the one for the glnKp is sequestered in a double-stranded hairpin structure. To provide additional support for this assertion, we switched the 5’-3’ order of the *mCherry* and *GST* genes in our circuit (Fig. 2C). In the new construct only the *GST* gene was subject to insulation, while mCherry was not subject to any putative inhibition encoded in its RNA sequence. We then measured the level of GST RNA, mCherry expression, and GST protein levels. The plot shows that while the GST RNA level remained approximately the same for both the insulated and non-insulated configurations (Fig. 2C-left), the amount of GST protein was sharply reduced when the gene was placed downstream of the insulator element within the glnKp promoter (Fig. 2C-right) in a manner similar to the reduction in mCherry fluorescence (Fig. 2B). Consequently, insulation functions independently of downstream genetic context, consistent with it being encoded as an independent regulatory element at the RNA level.

### Oligo library (OL) analysis of glnKp mutants

To further examine our hypotheses, to explore and characterize the insulating sequence and mechanism encoded by glnKp, and to check for its prevalence in other bacterial genomes, we constructed an oligo library (OL) of 12758 150-bp variants (Fig. 3A). The OL was synthesized by Twist Bioscience (for technical characteristics see Fig. S3 and Fig. S4) and inserted into the synthetic enhancer backbone following the method introduced by (Sharon et al., 2012). The OL was designed to screen both known σ^54^ promoters from various organisms and σ^54^-like sequences from genomic regions in *E. coli* and *V. cholera* and examine the silencing effect, essentially searching for similarities to the phenomenon observed in glnKp and for their potential sequence determinants. In addition, the OL was designed to characterize 134 glnKp mutational variants. Finally, the OL was designed to conduct a broader study of the contextual regulatory effects induced by a downstream genomic sequence, in either a sense or anti-sense orientation, on an active upstream promoter positioned nearby. Each variant consisted of a pLac/Ara promoter, followed by a variable sequence, an identical RBS, and an mCherry reporter gene, thus encoding a 5'UTR region with a variable 50 bp region positioned at +50 bp from the pLac/Ara TSS (Fig 3A). Similar to the experiment shown in Figure 2, each plasmid also contained an eYFP control gene to eliminate effects related to copy number differences and to enable proper normalization of expression values. Sort-Seq combines the OL with fluorescence-activated cell sorting and next-generation sequencing (Sharon et al., 2012), yielding the distribution of e.l.r for each sequence variant. Figure 3B shows the e.l.r distribution profiles for 10438 variants with sufficient total number of sequence counts (n>10, see Materials and Methods for details), revealing a broad range of distributions of expression levels. While a significant percentage of the variants showed low mean e.l.r, similar to what was observed for glnKp, a non-negligible set of variants produced high and intermediate expression levels (Fig. S5 top and bottom show representative examples), indicating that a combination of regulatory mechanisms encoded within these sequences may underlie this distribution of expression levels.

**Figure 3:**
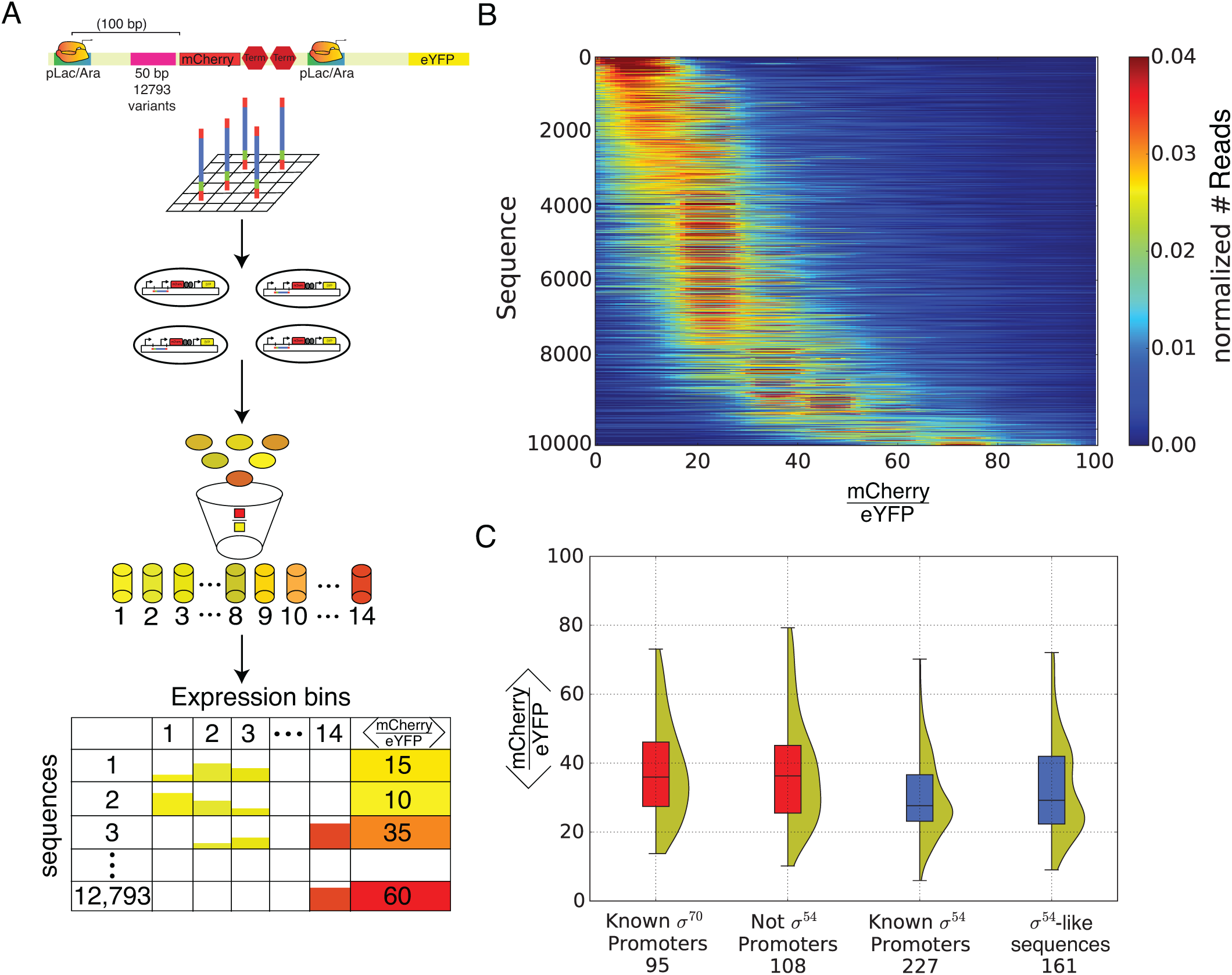
Oligo-library analysis of the insulation phenomenon. (A) Oligo library design and schematic description of the protocol. In brief, the synthesized oligo library (Twist Bioscience) was cloned into *E. coli* competent cells which were then grown and sorted by FACS into 14 expression bins according to mCherry to eYFP fluorescence or expression level ratio (e.l.r). DNA from the cells of each bins was barcoded and pooled into a single sequencing run to produce an e.l.r profile for each variant. For details see Materials and Methods. (B) Library expression distribution. Heat map of smoothed, normalized number of reads per expression bin obtained for 10438 analyzed variants ordered according to increasing mean e.l.r. (C) Violin plots showing mean expression value distribution for the following variant groups in the library: known σ^70^ promoters (95 variants), no promoter (108 variants), known σ^54^ promoters (227 variants), and σ^54^-like sequence (161 variants). σ^54^-like sequence variants were selected based similarity to the core promoter consensus sequence(Barrios et al., 1999) Associated with Figure S3, S4, and S5, Table S4.

To examine the observed silencing and the hypothesized insulation in broader context beyond glnKp, we analyzed the mean e.l.r values for four groups of variants in our library: annotated σ^54^ promoters from various organisms obtained from the list compiled by(Barrios et al., 1999) and EcoCyc database, annotated σ^70^ promoters obtained from EcoCyc, σ^54^-like sequences (score >10 – see method details), and non-σ^54^-promoters who were scored >0.9 and <0.5 by our consensus σ^54^ -promoter calculator respectively (see Materials and Methods). Checking the distributions of mean e.l.r within defined classes of variants (Fig. 3C), we observed that the "not σ^54^ promoter" class and the annotated σ^70^ distributions are shifted towards higher expression levels as compared with the annotated σ^54^ promoters and σ^54^-like distributions. A non-parametric Mann-Whitney U-test shows that the mean e.l.r computed for each of the two former classes is significantly higher than those of the latter classes (p-value < 10^−4^ compared to annotated σ^54^ promoters and p-value < 0.01 compared to σ^54^-like variants). This analysis suggests that there may be a conserved sequence determinant within the σ^54^-variant classes that contributes to this shift-down in expression levels.

### Oligo library (OL) analysis of broader insulation

To search for potential sequence determinants that may be associated with the differences in e.l.r distributions, we performed a DRIMust k-mer search on the OL variants sorted by mean e.l.r. values. DRIMust is a tool designed to identify enriched sequence motifs in a ranked list of sequences (Eden et al., 2007; Leibovich and Yakhini, 2012; Leibovich et al., 2013). Our analysis revealed that a single CT-rich consensus motif is enriched in the silenced variants (p-value < 10^−54^, minimum hypergeometric (mHG) test). We plot the results for 388 variants that are either annotated σ^54^ –promoters (227 variants) or σ^54^-like sequences (score >10 – 161 variants) in Fig. 4A. The consensus motif is derived from a list of ten 5 bp CT-rich features, each enriched in the top of the ranked list (see Fig. 4A-top-right). We call these enriched 5-mers “E5mers”. In the left panel, we plot the mean e.l.r value for each variant, ordered by decreasing e.l.r from top to bottom. The middle panel of Fig 4A shows the position of each E5mer, marked by a brown line in the corresponding variant on all 388 putative and annotated σ^54^ –promoters. In the right panel, we show a 20 variant running average of the number of E5mer occurrences. This analysis shows the correlation between the presence of an E5mer and the mean e.l.r value in the σ^54^ –promoter group. Together, the plots show that a high concentration of CT-rich motifs close to the purine-rich sequence that encodes the Shine-Dalgarno motif (SD – positioned at +17) is strongly associated with variants leading to low mCherry to eYFP fluorescence ratio.

**Figure 4:**
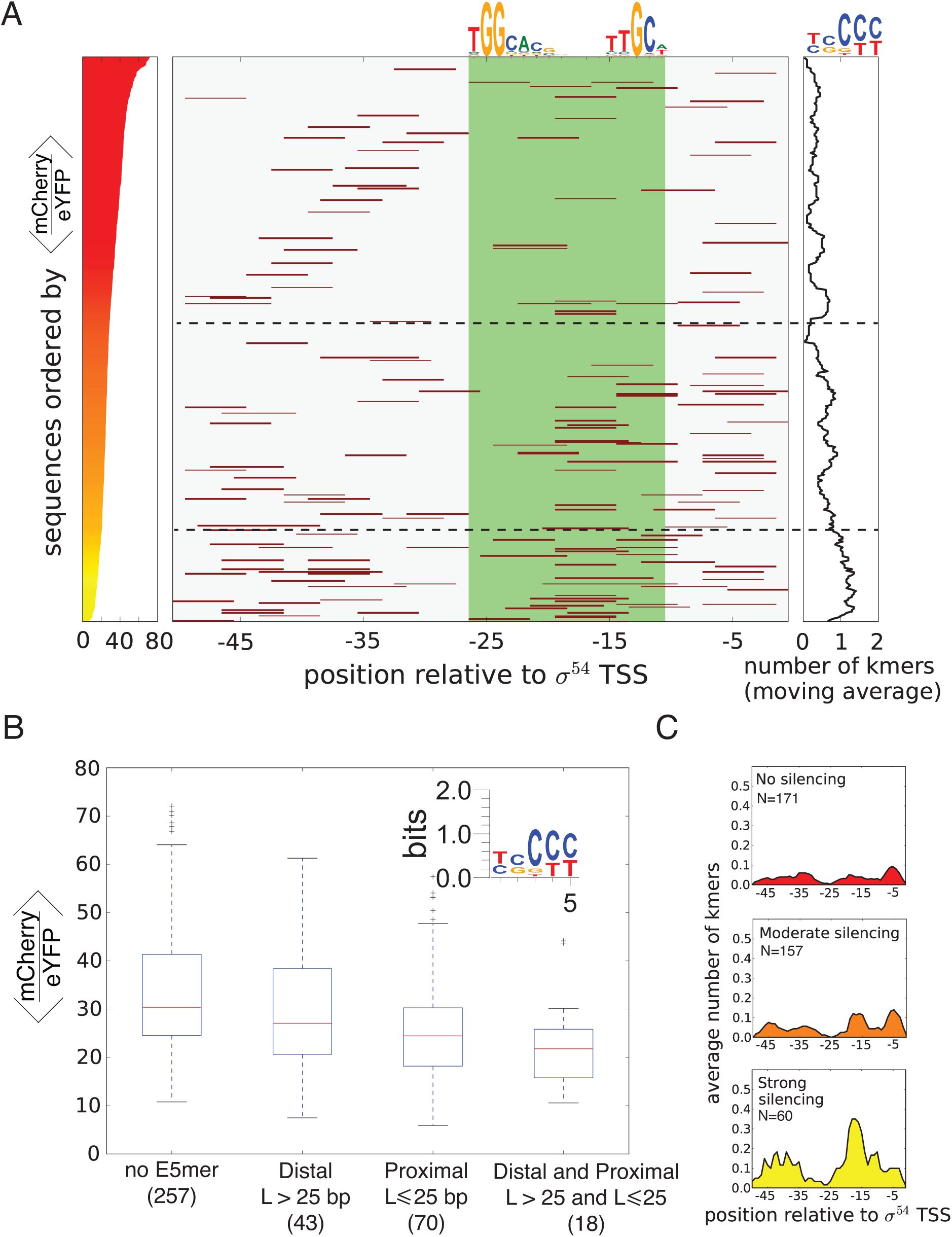
Insulation phenomenon is prevalent in other σ^54^ promoters. (A) Left: heat map ordering of the examined variants by mean e.l.r value, with silenced variants at the top. Middle: for each variant in the left panel, each enriched 5mer (E5mer) appearance is marked by a brown line at its position within the variant sequence. (Green shade) σ^54^ core promoter region. Top: σ^54^ core promoter consensus sequence(Barrios et al., 1999) Right: Running average on the number of E5mers observed within a variant in the ordered heat map. Top: a PSSM summarizing a multiple alignment of the E5mers found with DRIMust. (B) Box-plot showing groups of σ^54^ –like and annotated σ^54^ promoters differentiated by the location of 5Emers within the 50bp variant sequence. (C) Plots depicting the average number of E5mers found per position on the 50 bp variants for putative and annotated σ^54^ promoters. Variants are grouped by strong (bottom panel and variants below lower dashed line in panel B), moderate (middle panel and variants in between dashed lines in panel B), and no (top panel and variants above middle dashed line in panel B) silencing variants respectively. Associated with Table S5 and S6.

To further examine the dependence of the mean e.l.r on the position of E5mers within the annotated and putative σ^54^ promoters, we grouped the 388 sequences into four classes (Fig. 4B): containing no E5mers, E5mers located at a distal position from the σ^54^ promoter’s TSS (i.e. more than 25 bp upstream of the TSS), E5mers located at a TSS proximal position (i.e. less than 25 bp), and E5mers located at both the proximal and distal positions. In addition, we plot in Fig. 4C the average number of E5mers per position (with the σ^54^ 10 promoter TSS defined as 0) depicted over three regimes: strong insulation (mean e.l.r<20), moderate insulation (20<mean e.l.r <30), and no/weak insulation (mean e.l.r>30). We chose 25bp as the threshold due to the presence of a conserved sequence in the σ^54^ core promoter region (Fig. 4B-center-top) that is not an enriched region. The data shows (Fig. 4B-C) that there is a clear correlation between the number and position of the E5mers to the mean e.l.r. In particular, an ANOVA (See Tables S5 and S6 test on the mean e.l.r and the position of the E5mers shows that both proximal and distal E5mers affect the mean e.l.r, with the proximal effect being much more significant than the distal effect (p-value < 10^−6^ and 0.05 respectively). The test also shows that these effects are additive. The pattern of insulation with E5mers at distal and proximal locations was detected in 18 annotated and σ^54^-like promoters from various organisms (see list in SI), and is consistent with the observations for the glnKp, which also manifested this pattern. Overall, we found 60 strongly insulating σ^54^ promoters and σ^54^-like sequences (e.l.r <20, Fig. 4D-bottom) out of the 388 tested in our OL.

### Deletion of CT-rich motifs relieves the insulation effect

To provide further evidence that the CT-rich segments identified in our library underlie the insulation phenomenon, we analyzed 123 mutant variants of the glnK promoter that yielded *n>10* sequence counts (Fig. 5A). The figure shows the original glnKp sequence context at the top, and the set of mutations to the original context for each mutant plotted as individual lines of text over a red shaded box below. The lines of text over-lay a colored map corresponding to previously identified regions in the glnKp context. These regions include the CT-rich flanking regions (blue), core sequence of σ^54^ promoter (light green), and unclassified flanking region (green). We arranged the mutant glnKp variants in order of increasing e.l.r value from top to bottom with the most silenced variant at the top line (see heat-map gradient on the right, corresponding to the e.l.r value measured for each glnKp mutant variant). The figure shows that the mutations in the core sequence region and in the distal CT-rich region (left - blue shaded region) did not correlate with increasing e.l.r, but are rather evenly distributed throughout the mean e.l.r. range. However, increased amount of mutations in the proximal CT-rich segments of the flanking region (right - blue shade) and in positions immediately upstream of the TSS correlate strongly with elevated mean e.l.r. In particular, mutations in the 7 nucleotide CT-rich region (centered at -4) into a G or an A yielded the largest increase suggesting that the insulating effect was likely abolished.

**Figure 5:**
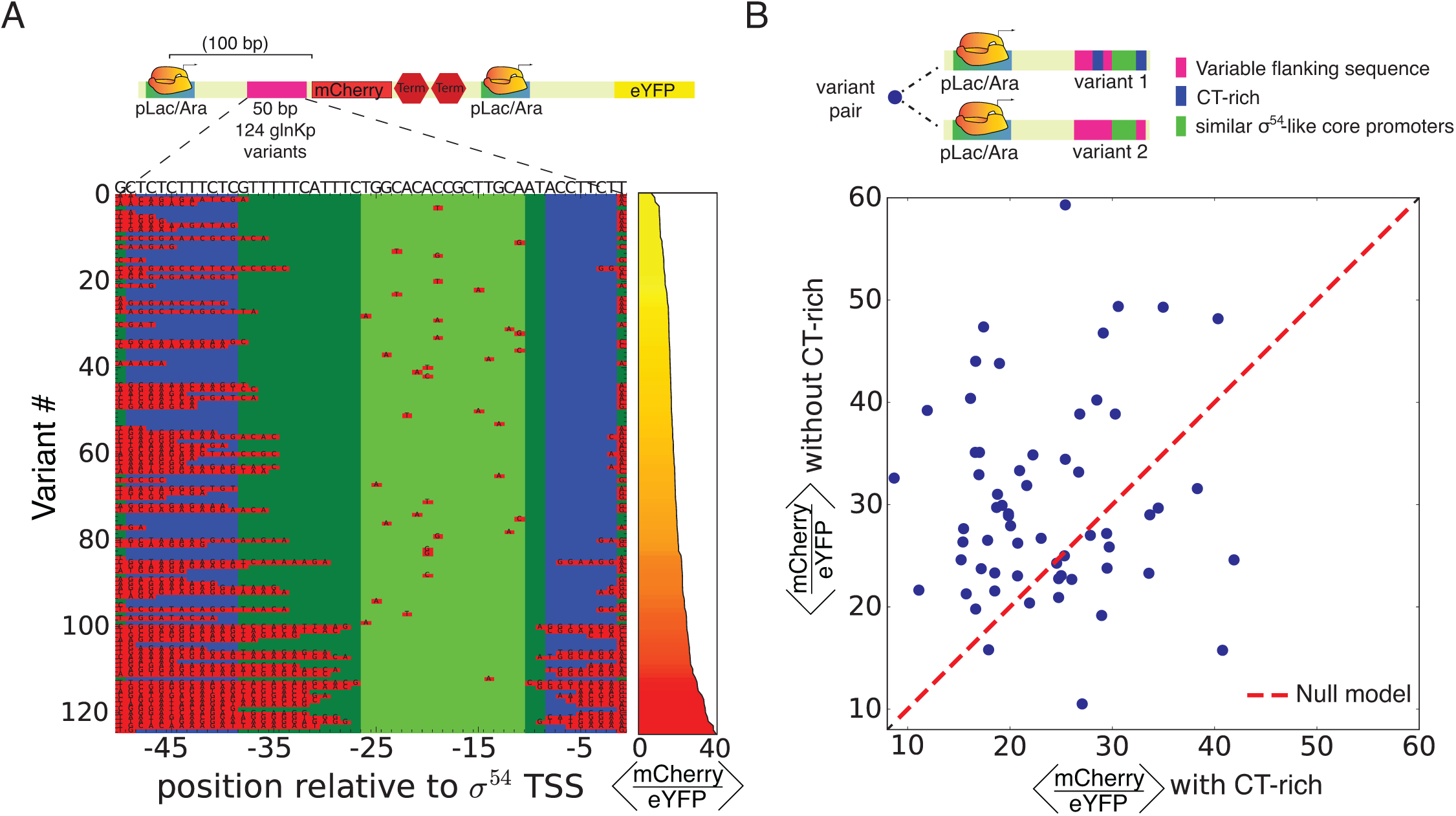
Analysis of CT-rich deletions in insulated variants. (A) Analysis of the glnKp mutation subset of the library. (Top) circuit schematic showing the location of the variant sequence. (Bottom) glnKp mutated variants with glnKp sequence shown on top. Flanking regions, core σ^54^ promoter, the CT-rich regions, and mutations are denoted by dark green, light green, blue, and red boxes, respectively. Right panel denotes the mean e.l.r value using a yellow-to-red scale. (B) (Top) schematic showing the variant-pair contents by region: (pink) variable flanking sequence, (blue) E5mers, (green) core σ^54^ promoter. (Scatter plot) Analysis of expression levels of σ^54^-promoter-like sequence pairs with and without a CT-rich region. Each point represents a pair of nearly-identical sequences (see methods), one containing a CT-rich region in the flanking region and the other does not. In most pairs, the CT-rich containing sequence (X axis) presents lower e.l.r values (yellow points) than the one lacking a CT-rich region (Y axis) indicating that the CT-rich region is related to the silencing phenomenon.

We next investigated pairs of variants with high sequence similarity in the putative σ^54^ core promoter region to further compare the effect of E5mers presence on insulation. Variants were considered to be similar if at least 11 of the 16 core-promoter bases were identical. Each pair consisted of one sequence variant whose flanking region contained the E5mer identified in Fig. 4A, while the other did not. Furthermore, the remaining 29 bps of unclassified flanking region were identical in at least 21 of the 29 bases (See Fig. 5B-top for a schematic description of the variant pairs). Thus, this analysis compares σ^54^-promoter-like variant that contains an E5mer to a close one that lacks it. In the plot (Fig 5B-bottom), each variant pair is represented by a circle whose x-coordinate and y-coordinate correspond to the mean e.l.r value measured for the variant that contains and lacks the E5mer, respectively. To check if there is a bias towards higher e.l.r values for variants that lack the E5mer, we add to the scatter plot a dashed line representing the NULL assumption that *x=y*. The plot shows in comparison to the dashed line the scatter plot is biased towards higher e.l.r values, indicating that variants that have higher e.l.r values (as compared to their paired variant) are typically those lacking the E5mer. Over-all we found 72 non glnKp σ^54^-core-promoter-like variant pairs in our library with 52 pairs positioned above the *x=y* line and 20 below, yielding a p-value of <= 6.2x10^−7^ (two-tailed t-test vs an equal means null hypothesis). This analysis further supports the necessary and sufficient silencing role of the E5mers in our OL.

### CT-rich segments are depleted in bacterial genomes upstream of putative RBS

Next, we performed bioinformatics analysis to examine the occurrence of aSD:SD potential interactions around σ^54^ promoters in the native *E. coli* genomic context. To this end, we counted proximal occurrences of aSD:SD sequences around σ^54^ promoters and compared the numbers to such occurrences around all other promoters (i.e. σ^70^–like). We first identified N=8174 putative promoters and their respective TSSs (Salgado et al., 2013), B=91 of which are annotated as σ^54^ promoters. We then identified the first putative Shine-Dalgarno (purine-rich) hexamer downstream of the TSS, and the best matching (Hamming) aSD (pyrimidine rich) hexamer upstream of the TSS, both up to 50bp (Fig. 6A). In a total of B=1383 of the N=8174 promoters we found a potential near-perfect (Hamming distance ≤ 1) proximal aSD:SD pair (See Fig. 6B), b=25 of them in the n=91 σ^54^ promoters. Under a Hypergeometric model this yields an enrichment at p-value<0.008 for the occurrence of these sequences within σ^54^ promoters.

**Figure 6:**
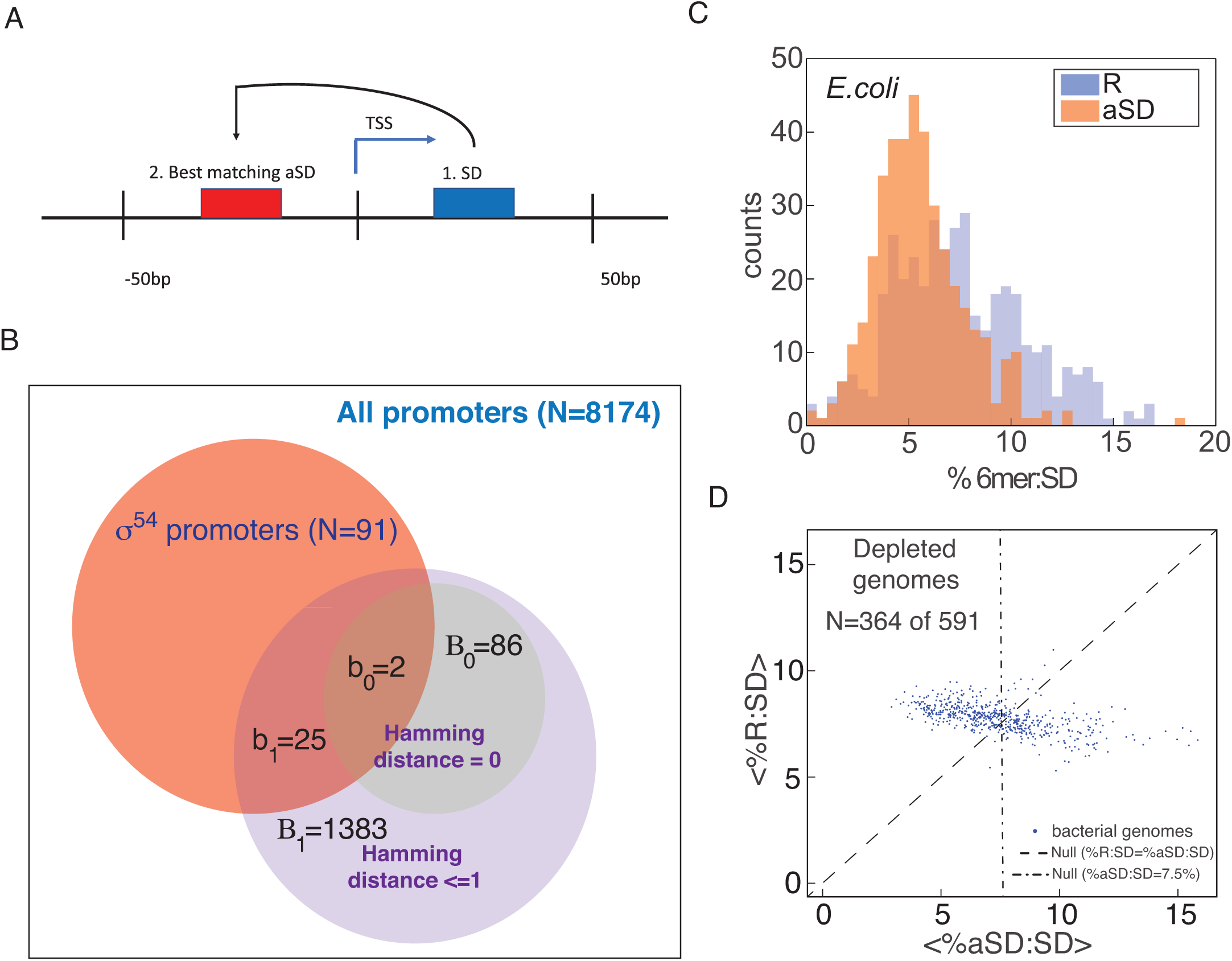
Analysis of prevalence of CT-rich k-mers around *E.coli* promoters and in other bacterial genomes. (A) A scheme for the analysis of the occurrences of aSD:SD around σ^54^ TSS positions. 1. Step i - we scan the 50bp region downstream to TSS and locate the SD. 2. Step ii - the best matched aSD, i.e. the hexamer which is the best Hamming reverse complement to the SD we found in Step i, is found in the 50bp upstream the TSS. (B) Venn diagram for promoters found with an aSD sequence that is either a perfect match or at most 1bp away. Square: the space of all putative *E. coli* promoters. Red circle: the space of all putative *E. coli* σ^54^ promoters. Green and purple circles: promoters that possess either a perfect aSD match or one off by 1bp (C) Distribution of % proximal occurrences (%6mer:SD) within 300bp separation (see Methods detail) of CT-rich to GA-rich (aSD:SD) pairs (orange) as compared with the % proximal occurrences of random to GA-rich (R:SD) hexamer pairs for *E. coli*. (D) Scatter plot for 591 mesophile and psychrophile bacterial genomes, where each genome is represented by the mean value for the aSD:SD (x-axis) and R:SD (y-axis) % proximal occurrence distributions. Dashed line (x=y) corresponds to the null model assuming that mean aSD:SD should equal to mean R:SD. Vertical dashed-dot line (x=7.5% occurrences at 300bp) corresponds to the null expected value. Associated with Figure S6 and Table S7.

To put this enrichment of aSD:SD occurrence near σ^54^ promoters in an even broader bacterial context and also to check for potential promoter sequence bias which may affect this finding, we needed to carry out whole genome analysis on multiple bacterial genomes. We hypothesized that if pyrimidine-rich hexamers encode a regulatory function then the prevalence of *aSD:SD* will be reduced as compared to *random:SD* occurrences, where *random* is some random hexamer. In order to test this hypothesis, we developed a specialized algorithm to quantitatively assess the prevalence of *aSD* and *random (R)* motifs upstream of putative SD hexamers at the genomic level (see method details and Fig. S6 for detailed description of algorithm). In brief, we first constructed a list of all hexamers (4096) and for each one of them we calculated the free energy value (ΔG) for its hybridization with the 16S rRNA of *E.coli* (as in (Li et al., 2012)). We ranked this list of hexamers from low to high free energy value, which corresponds to higher to lower probability for hybridization with the 16S rRNA. The top 20 hexamers were defined as the best 20 putative SD hexamers. Next, we computed a set of 20 pyrimidine-rich hexamers with the highest PSSM scores according to our aSD E5mer motif (see Fig. 4A top right for logo). Finally, we computed a set of 20 randomized hexamers, which did not score highly for either motif (See Table S7 for all hexamers). To compare the occurrence frequency of proximal *aSD:SD* to that of *R:SD* (*random:SD)*, we first identified all of the hits in the genome for a particular SD hexamer. An *occurrence* is then defined as the location of the first appearance of a particular aSD hexamer upstream of one of the hits. Next, we computed all occurrences in the genome for a particular *aSD:SD* and *R:SD* pairs in the range *10<d<10000,* where *d* is the distance in base-pairs upstream of the putative purine-rich hexamer. Finally, we computed the percentage of proximal pairs (*%aSD:SD* - defined as the percentage of occurrences that fall in the range *10<d<300*) for each pair of *aSD:SD* and *R:SD* hexamers. This process yielded a distribution of 400 percentage numbers for both randomized (R) and CT-rich (aSD) to GA-rich (SD) segments. As an example, we plot the distribution of percentage of proximal pairs obtained for *E. coli* in Fig. 6C. The plot shows that in *E. coli aSD:SD* proximal occurrences (orange) are significantly depleted as compared to the *R:SD* (Wilcoxon p-value < 3.83×10^−12^).

We applied our aSD:SD prevalence algorithm to 591 psychrophile and mesophile genomes (i.e. bacteria that thrive in the temperature range of -10-40°C), obtaining two percentage distributions (as in Fig. 6C) for each genome. We then computed the mean for each distribution, allowing us to represent each genome by these two numbers. To quantitatively assess whether the depletion of aSD:SD is prevalent in mesophilic genomes, and not just specifically in *E. coli*, we represent each genome as circle in a scatter plot with the x- and y-coordinates set as the mean proximal *R:SD* and *aSD:SD* occurrences respectively. We found that while the value for the mean percentage *R:SD* for all genomes is scattered tightly around the null expected 7.5%, the aSD:SD genomic scatter varies widely over a considerably larger range. In addition, assuming no significant regulatory function for the aSD motifs, we should expect to find approximately half the genomes above the dashed line (x=y) and half below. Instead, a disproportionately large number of genomes (364 of 591 or ~62%) are found with a larger mean R:SD percentage than aSD:SD. Finally, 342 of 591 (or 57.87%) of the genomes are found with mean aSD:SD < 7.5% (left of the vertical dashed-dotted line). In summary, while merely a statistical finding that may also reflect other evolutionary constraints (e.g. depletion of GGGG due to selection against G-quadruplex formation, or possible codon selection in the 3’terminus of genes), the enrichment of aSD:SD pairs in the vicinity σ^54^ promoters as compared with other promoter types, together with the over-all depletion of aSD:SD proximal occurrences in bacterial genomes, is consistent with a natural role for the insulating mechanism we observed in the synthetic systems. One that is mostly utilized within the context of σ^54^ promoters as defined by the annotation used herein.

### Modeling insulation can help improve performance of gene expression algorithms

Finally, our study has consequences for synthetic biology applications, where circuit design often fails due to “context-related” effects. Here, well-characterized parts, which can originate from different organisms, are often tailored together. Our results imply that careful attention needs to be paid to the flanking or bridging sequences, which are used to stitch the parts together, as they may encode unwanted regulatory effects. To get another perspective on the regulatory effect generated by “context” or “flanking” regions, we first used RNAfold (Hofacker et al., 1994) to compute the probability for the RBS to be sequestered in a secondary structure for each variant in our library (we only used the 5’UTR region of each variant for this computed). We then plotted the mean e.l.r for each variant as a function of the computed probability for the RBS to be sequestered (Fig. 7) showing a trend of increasing e.l.r values as a function of decreasing probability for RBS sequestration. Next, we computed the predicted expression level of our OL variants using RBSDesigner (Na and Lee, 2010). This is a tool designed to predict protein level for a given sequence based on RBS availability and affinity to the ribosome. In Fig. 7A we plot in blue the binned distribution of the RBS designer predictions as a function of the probability of the same variants to have their RBS sequestered. The data shows that the RBS designer predictions for our sequence variants are widely distributed and uncorrelated with the structural prediction generated by RNA fold. By comparing these predictions to the increasing trend observed with the experimental data (red bins), it seems the RBS Designer tool does not take into account some important regulatory phenomenon.

**Figure 7:**
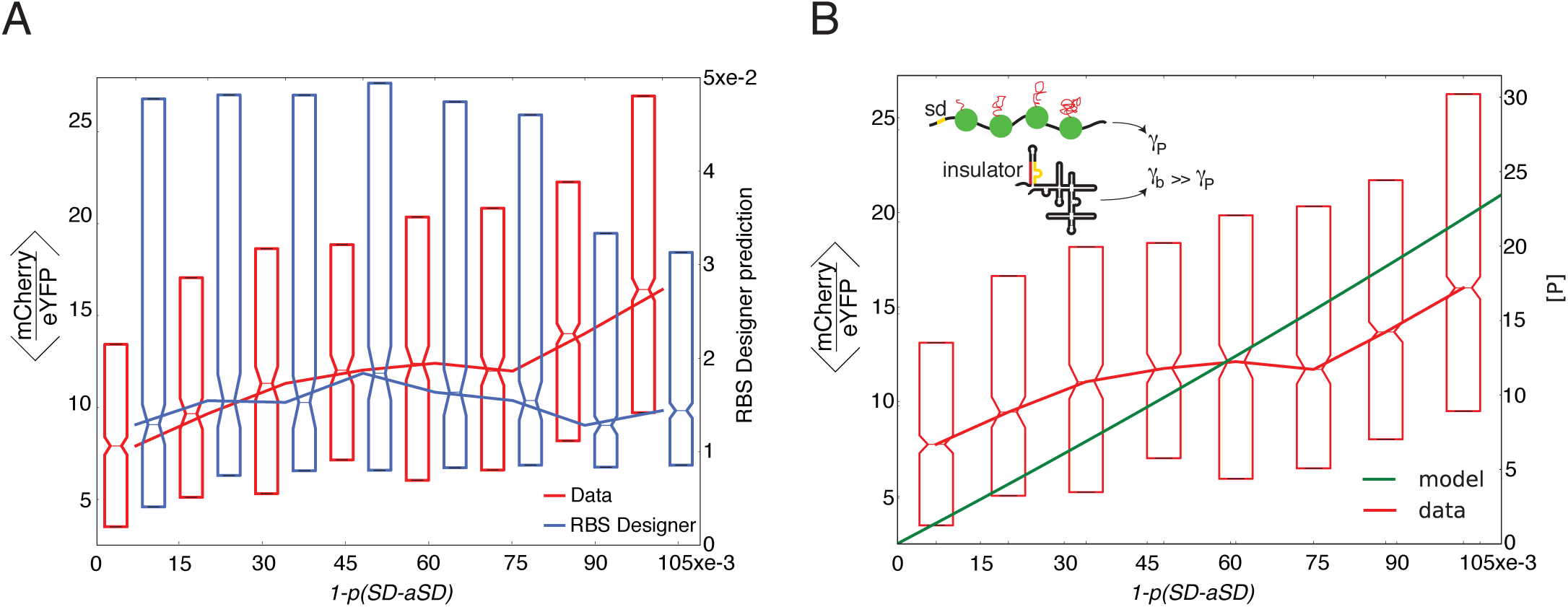
Comparison of RBS designer tool and Insulation model for predicting gene expression on our library. (A) Binned mean fluorescence ratio of the library variants plotted in box-plot form according to the probability of the RBS to be unbound (red) together with the RBS Designer tool predicted mean expression levels for each variant sequence (blue). The comparison shows that the RBS Designer predicted RBS availability does not reflect the measured mean fluorescence ratio, and as a result is a poor predictor for our data. (B) The library mean fluorescence ratio plotted in box-plot form as a function of the probability of the RBS to be unbound (red). The prediction from the degradation model is plotted for comparison (green). (Inset) A scheme of the degradation model (see methods detail). Top: the translated pearled phase with low degradation rate. Bottom: the non-translated branched phase with high degradation rate.

To qualitatively capture the experimental trends, we devised a simple gene expression model that takes into account the probability that the RBS is sequestered. We assumed different degradation rates for RBS sequestered and non-sequestered mRNA structures (see Fig. 7B – inset and detailed methods). Finally, we used realistic constant rates for other kinetic processes (e.g. transcription rate, translation rate, etc. (Milo & Philips, 2016)). In Fig. 7B we plot as bins the measured mean e.l.r as a function of the RNAfold computed probability for RBS to be sequestered in a secondary structure as a result of aSD:SD interaction. In addition, we also plot the predicted expression level ([P] - right y-axis) from our two-rate degradation model as a function of the aSD-SD secondary structure probability (green-line). The plot shows that the model qualitatively tracks the trends in the experimental data exhibiting a mild increasing slope in “predicted” expression level as a function of decreasing probability for RBS sequestration. Thus, taking into account the insulation phenomenon that we describe in this work can improve the performance of gene-expression prediction tools.

## Discussion

We used a synthetic oligonucleotide-library (OL)-based approach, following protocols introduced in recent years (Kinney et al., 2010; Sharon et al., 2012), to uncover and study a context-dependent phenomenon of translational insulation in σ^54^ promoters. To do so, we designed and constructed a library of over 10000 closely-related transcriptional read-through circuits, with σ^54^-like variants serving as the variable sequence. The insulation phenomenon was characterized for a subset of annotated σ^54^ promoters, and additionally for σ^54^-like sequences in *E. coli* and *V. cholera*, which may be either unknown promoters or σ^54^ intergenic binding sites. We also carried out an extensive mutational analysis using the OL on the *E. coli glnK* promoter to determine the precise sequence element responsible for the insulation phenomenon in glnKp.

Insulation was found to be dependent on the prevalence of short (3-5 nucleotide) CT-rich sequences, which were distributed at various positions within the 50 bp variants. Strength of insulation depended on the number and proximity of CT-rich segments to the putative σ^54^ promoter TSS. Since these CT-rich segments cannot be transcribed by the σ^54^ promoter to which they belong, insulation seems likely to be associated with transcriptional read-through, when the polymerase originates from another locus upstream. Together, the segments create a cumulative effect that possibly triggers a collapse of the RNA molecule into a “branched phase” via aSD:SD interaction (Schwab and Bruinsma, 2009) due to the lack of translocating ribosomes on the mRNA molecule leading to rapid degradation of the insulated mRNA molecule (Richards et al., 2012). This hypothesized mechanism is supported by modelling evidence as well as by comparing variants that mostly differ by the presence of the putative aSD element. In this work, this comparison includes a high coverage analysis for glnKp and a sparser analysis for other variants. However, there are likely other “context” related regulatory mechanisms encoded into these variants that may contribute to the wide distribution observed for the OL gene expression profile. Those remain obscure due to the unavoidable partial coverage provided by our library.

Previous studies have implicated aSD-SD interactions, and other secondary structure formations involving the RBS in a variety of regulatory phenomena. These include riboswitches, up-regulation via RNA binding protein interactions with RNA (Babitzke et al., 2009; Winkler and Breaker, 2005), modulating expression levels with partially stable structures (De Smit and Duin, 1990; Schwartz et al., 1981), and inducing translation via S1-interaction (non-SD initiation (Komarova et al., 2002)). Other studies (see (Nakamoto, 2006) for review and references therein) have suggested that translation initiation is inhibited when the AUG is sequestered in a double stranded structure, and that this can be avoided by either having a non-structured 5’UTR region (Scharff et al., 2011), or via an accessible Shine-Dalgarno (SD) sequence. Therefore, if the SD sequence is also sequestered, then the likelihood of AUG inaccessibility to translation initiation will be high. Finally, recent work argued for the presence of a particular group of sequences upstream of the RBS, which play a role in recruiting the small subunit of the ribosome possibly by destabilizing secondary structures (Campo et al., 2015). The insulation phenomenon described in the present work and its relative prevalence in annotated σ^54^ promoters is another regulatory manifestation of the effects of RNA secondary structure, and of the effects of aSD:SD interactions in bacteria.

Given the potency of this regulatory effect and its statistical depletion in most bacterial genomes, why is there an enrichment in the sub-class of σ^54^ promoters? Unlike most bacterial sigma factors that are members of the of σ^70^ family and encode a niche response, σ^54^ promoters are unique. The polymerase is unable to initiate transcription by itself, but rather absolutely requires the energy of ATP hydrolysis via the binding of an associated bacterial enhancer binding protein. As a result, σ^54^ promoters do not suffer from promoter leakage and are usually fully repressed when the bacterial enhancer binding protein is absent. The encoding of the aSD sequences in the non-transcribed portion of these promoters generates another level of security against errant transcriptional events, ensuring that σ^54^ -regulated genes are not produced when there is accidental transcriptional read-through from an upstream promoter. Recent analysis has revealed a common functional theme across multiple bacterial species, which can provide an explanation for the additional measures used in these promoters against leaky or errant transcription. The analysis of (Francke et al., 2011) has shown that σ^54^ promoters predominantly regulate genes that control the transport and biosynthesis of the molecules that constitute the bacterial exterior, thus affecting cell structure, developmental phase, and interaction potential with the environment. For instance, in *M. xanthus*, there are many σ^54^ promoters that have been associated with fruiting body development (Jakobsen et al., 2004). Thus, it is possible that the presence of CT-rich insulators encoded within some σ^54^ promoters may be attributed to preventing metabolic and developmental consequences, which only in rare circumstances are needed for survival.

While our results provide additional support to the observations that short CT-rich sequences in the vicinity of an RBS can affect expression, they are by no means the only "context" dependent effect, which can be encoded in an accidental fashion, in the design of synthetic circuits. To avoid other such context-dependent regulatory phenomena, we believe that it can be useful to apply OL based approaches (reviewed in (Peterman and Levine, 2016)) as exemplified here. This experimental approach constitutes a reliable systematic methodology in synthetic biology to support gene circuit design and construction. It is a high-throughput controlled mutagenesis that can serve as “debugging” analysis. Sort-seq approaches can be utilized to identify, characterize, and filter out unknown context-related effects, facilitating a desirable outcome without resorting to an iterative design/characterization process. Studies based on Sort-seq can therefore lead to improved performance of design tools, such as discussed in comparing RBS-designer to a refined model in Fig 7. In summary, our findings advance our understanding of translational regulation in bacteria. From a practical perspective, our findings provide potentially useful refinements of guidelines to be used in synthetic biology designs.

## Supplemental Information

6 Supplementary Figures

7 Supplementary Tables

2 Supplementary Data sets

## Acknowledgments

This project received funding from the European Union's Horizon 2020 Research And Innovation Programme under grant agreement no. 664918 - MRG-GRammar, the Israel Science Foundation through Grant No. 1677/12, the I-CORE Program of the Planning and Budgeting Committee and the Israel Science Foundation (Grant No. 152/11), and Marie Curie Reintegration Grant No. PCIG11-GA-2012-321675. The authors would like to acknowledge the Technion's LS&E staff (Tal Katz-Ezov, Efrat Barak, Anastasia Diviatis) for help with sequencing and FACS, and Adina Weinberger, Yitzhak Pilpel, Roy Kishony, Shay Ben-Elazar, Noa Katz, Beate Kaufmann, Yaroslav Pollak, and Inbal Vaknin for useful discussions.

## Author contribution

LL designed and carried out the experiments for both the initial σ^54^ and OL experiment. LA designed and carried out data analysis for the OL library results and modeled the data. OS designed and performed the bioinformatics genomic context analysis. RC, SO, and SG designed and carried out the Δ*rpoN* and GST experiments. OA assisted with some of the experiments. RA and ZY supervised the study. RA, ZY, SG, LL, LA, and OS wrote the manuscript.

## Supplemental Information

## Method Details

### Synthetic enhancer construction

Synthetic enhancer cassettes were ordered as dsDNA minigenes from Gen9, Inc. each minigene was ~500 bp long, and contained the following parts: NdeI restriction site, variable UAS, variable σ54 promoter, and KpnI restriction site at the 3’ end. The UAS and σ54 promoter were separated by a looping segment of 70 bp. For sequence details see Supplementary Note 1 and Supplementary Tables 1 and 2. Insertion of minigene cassettes into the plasmid was done by double digestion of both cassettes and plasmids with NdeI and KpnI, followed by ligation to a backbone plasmid containing an NtrC switch with TetR binding sites24 and transformation into 3.300LG E. coli cells containing an auxiliary plasmid overexpressing TetR. Cloning was verified by Sanger sequencing.

### Synthetic enhancer fluorescence measurement

Starters of strains containing the enhancer plasmids were growth in LB medium with regular antibiotics overnight (16 hrs). The next morning, the cultures were diluted 1:100 into fresh LB and antibiotics and grown to OD600 of 0.6. Cells were then pelleted and medium exchanged for BA with antibiotics. Fluorescence was measured after an additional 2 hrs of growth in BA. Measurements of mCherry and eYFP fluorescence were performed on a FACS Aria IIIu (without sorting).

### NtrC switch design

The NtrC switch was based on positive-feedback synthetic enhancers that were described previously (Amit et al., 2011; Brunwasser-Meirom et al., 2016). In brief, a UAS containing the glnAp1 promoter was positioned 121 bp upstream from the transcriptional start site of glnAp2 and a *glnG* gene (which codes for NtrC) (see Fig. SS1A). In the intervening looping region, three TetR binding sites were placed with intersite spacing of 33 bp (“in phase”). TetR was expressed from a separate high-copy plasmid via a constitutive promoter. When anhydrous tetracycline (aTc) is absent, TetR binds to its sites and the switch is turned off. When sufficient aTc is present, TetR is removed from the looping region and NtrC is expressed at high levels due to the positive feedback. In Supp. Fig. S1B, we plot the dose response for glnKp as a function of aTc showing the sharp change in expression, which is indicative of NtrC being turned on in the system.

### UAS and promoter definitions

For plasmid design, we chose 5 *σ*^54^ promoters and 1 no-promoter control, each with 10 composite UAS sequences and 1 no-UAS control, resulting in 66 constructs. The promoters were naturally occuring in *E. coli*. We chose 2 strong, 2 weak, and 1 intermediate strength promoters. Supplementary Table S1 lists the promoter names and sequences. Supplementary Table S2 lists the UAS sites used in the experiment. All synthetic enhancers were constructed with an identical 70 bp looping sequence between the UAS and *σ*^54^ promoter. The sequence, which we designed so it would not contain any putative UAS sites, known TF binding sites, or annotated promoters, was: GGCCGTTGAGAAAAGCCTGTCCCACTAGGTGGGCGTTCCGGCCTTACAGAGCGAATGGCGTAGTGCCGCA.

### TOP10:ΔrpoN strain construction

An E. coli TOP10:ΔrpoN strain was created in our lab following the protocol described in 48, using Addgene plasmids pCas (#62225) and pTarget:rpoN (based on Addgene plasmid #62226, with N20 target sequence 5’CCGTCCTTAAGCGGATCCAA3’), and a linear repair oligo constructed using overlap PCR containing the genomic sequences immediately upstream and downstream of the rpoN gene. After curing both plasmids, the genomic deletion was sequence-verified using Sanger sequencing of the rpoN genomic region. The lack of rpoN transcripts was further verified using qPCR with primers targeting rpoN.

### RNA extraction and reverse-transcription

Starters of E. coli TOP10 or TOP10:ΔrpoN containing the relevant constructs on plasmids were grown in LB medium with appropriate antibiotics overnight (16 hr). The next morning, the cultures were diluted 1:100 into fresh LB and antibiotics and grown to OD600 of 0.6. For each isolation, RNA was extracted from 1.5 ml of cell culture using standard protocols. Briefly, cells were lysed using Max Bacterial Enhancement Reagent followed by Trizol treatment (both from Life Technologies). Phase separation was performed using chloroform. RNA was precipitated from the aqueous phase using isopropanol and ethanol washes, and then resuspended in Rnase-free water. RNA quality was assessed by running 500 ng on 1% agarose gel. After extraction, RNA was subjected to DNase (Ambion/Life Technologies) and then reverse-transcribed using MultiScribe Reverse Transcriptase and random primer mix (Applied Biosystems/Life Technologies). RNA was isolated from 3 individual colonies for each construct.

### qPCR measurements

Primer pairs for mCherry, eYFP and GST genes, and normalizing gene idnT, were chosen using the Primer Express software, and BLASTed (NCBI) with respect to the E. coli K-12 substr. DH10B (taxid:316385) genome (which is similar to TOP10) to avoid off-target amplicons. qPCR was carried out on a QuantStudio 12K Flex (Applied Biosystems/Life Technologies) machine using SYBR-Green. 3 technical replicates were measured for each of the 3 biological replicates. A CT threshold of 0.2 Was chosen for all genes.

### Western blot analysis

Two strains: RBS+GST+glnKp+mCherry and glnKp+mCherry+RBS+GST were grown overnight in LB. Cells were briefly centrifuged and lysed with 4x Laemmli sample buffer (BIORAD, #161-0747). Equal volumes of lysates were loaded on a SDS-PAGE. Proteins were transferred to a Nitrocellulose membrane (Millipore), blocked in 5% fat-free milk for 1 hour at room temperature, and then incubated with the following primary antibodies diluted in 5% milk overnight at 4 °C: GST 1:1000 (Santa Cruz, sc138) and DNAk 1:10000(Abcam, ab73473). The HRP-conjugated secondary antibodies: Goat Anti-Mouse (LSbio, LS-C60680) was diluted at 1:25,000 in TBST. Immunodetection was performed using Amersham ECL Prime Western Blotting Detection Reagent (GE, RPN2232).

### σ54 consensus binding site scoring

The consensus probability matrix for *σ*^54^ binding (Appendix 1) was based on the compilation of 186 σ54 promoters (Table 3 in (Barrios et al., 1999)). The genomes of E.coli and V.cholera were scanned using a Matlab script that assigns a *σ*^54^ probability score to all possible 16 bp-long sequences, based on similarity to the consensus site. In brief, each base in the 16 bp sequence is given a value of 0-1 according to SI-Table 2. The values of all 16 bases are summed, and the total is normalized by first subtracting the lowest possible total (1.679) and then dividing by the difference between the highest possible total (11.2) minus the lowest possible total (1.679), resulting in a final score in the range [0,1] (shown in the equation below).

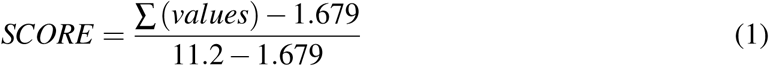

Genomic sequences with scores in the range [0.765, 1] were chosen as candidates for *σ*^54^ binding. Genomic sequences with scores in the range [0, 0.5] were chosen as candidates that were highly unlikely to bind *σ*^54^ and were taken as the “no promoter” group.

### OL design

Each variant included a unique 50 bp sequence, placed 120 bp downstream from the pLac/Ara promoter, and adjacent to an mCherry RBS, thus encoding a variable 5’UTR region with an interchangeable 50 bp region positioned at +50 from the TSS. The OL was designed to test both additional *σ*^54^ and putative *σ*^54^ promoters, from *E. coli* as well as other bacteria, for the silencing effect. In addition, we designed the OL to conduct a broader study of the contextual regulatory effects induced by a downstream promoter on an active upstream promoter positioned nearby in either a sense or anti-sense orientation. To do so, our library is composed of four sub-classes: a no-*σ*^54^ promoter set taken from *E. coli* and V*. cholera* genomic regions with low *σ*^54^ binding score (score<0.5), which was designed to form a non-coding positive control (130 variants). A set of 125 annotated *E. coli σ*^70^ promoters (devoid of any annotated TF binding sites) taken from the EcoCyc database (Keseler et al., 2013). A set of 275 annotated core *σ*^54^ promoters from multiple strains with their flanking sequences taken from both EcoCyc (Keseler et al., 2013) (98 variants) and (Barrios et al., 1999) (177 variants). A set of 134 mutant variants for the glnKp sequence in both the core elements and flanking sequences. 5715 variants with *σ*^54^ -like core regions mined from the *E. coli* and *V. cholera* genomes with a match score > 0.765 as compared with the *σ*^54^ consensus sequence (score =1, see eq. 1 above). Finally, all variants were encoded so they would appear in both sense and anti-sense orientations with respect to the pLac/Ara driver promoter. An OL of all variants was synthesized by Twist Bioscience. The library design contained 12762 unique sequences, each of length 145-148 bp. Each oligo contained the following parts: 5’ primer binding sequence, NdeI restriction site, specific 10 bp barcode, variable tested sequence, XmaI restriction site and 3’ primer binding sequence. The barcode and the promoter sequences were separated by a spacer segment of 23 bp (cassette design is shown in Fig. 1). We received 12758 sequences of the 12762 ordered.

### OL technical assessment

#### pre-next-generation sequencing

The library arrived as 974 ng of dried ssDNA and was diluted to 1 ng/*μ*l in TE. 1 *μ*l of the stock was examined using a high-sensitivity RNA kit on an Agilent Tapestation giving a clear peak of 150 bp. The library underwent PCR amplification to produce dsDNA products using Agilent Herculase II Fusion DNA polymerase (Catalog #600679) and the following PCR program: Step 1: 5 min – 95°. Step 2: 20 sec - 95°. Step 3: 20 sec - 50°. Step 4: 1 min - 68°. Step 5: 4 min - 68°. After calibration, an initial concentation 0.025 ng/*μ*l and 8 PCR cycles (final concentration of 21.4 ng/*μ*l) produced the best results in the cloning stage.

#### post-next generation sequencing

50 ng of the amplified library (8 cycles as described above) were sent to sequencing on an Illumina MiSeq (single run, 150 bp paired-ends) at the Technion’s Genome Center resulting in 4.3 million reads. Only read pairs in which there was 100% agreement on the sequence were used to reduce the effect of sequencing errors. This totaled in 1.7M reads that were aligned against the designed library to assess different properties as reported in Fig. S4.

##### Variant abundance

Based on 1.4M perfect match reads only, 99.27% (12665/12762) of the variants where detected with at least one read while 98.54% (12572/12762) where detected with at least 10 reads. The distribution of read counts per designed oligo has a mean of 110 reads and SD of 43. The distribution and its Lorenz curve are depicted in Fig. S4A-C. We studied the relationship between the sequence composition and the variant abundance, seeing a correlation coefficient of -0.54 as depicted in Fig. S4D.

##### Error rates

Each read was aligned to the library, using the variant with lowest edit distance as a match. All edit operations needed for match assignment were counted. These edit operations were used to assess the different synthesis error rates. Fig. S4E illustrates the calculation of the error rates. Each row represents a read, where blue reads are those with a perfect match to the designed variant. In reads that contain errors, the exact position of each error is marked, making it possible to generate nucleotide-specific analyses. Table S3 presents the different synthesis error rates by error type and base type.

Perfect match rate: 1 / 1.21 reads; 83%

Mismatch rate: 1 / 1550 bases

Deletion (1nt) rate: 1/ 1671 bases

Insertion (1nt) rate: 1/ 5133 bases

Deletion of length > 1: 1 / 1198 reads

Insertion of length > 1: 1 / 6402 reads

### OL cloning

Oligo library cloning was based on the cloning protocol developed by the Segal group (see Supplementary Note 2 for additional details). Briefly, the 12758-variant ssDNA library from Twist BioScience was amplified in a 96-well plate using PCR, purified, and merged into one tube. Following purification, dsDNA was cut using XmaI and NdeI and dsDNA with the desired length was gel-separated and cleaned. Resulting DNA fragments were ligated to the target plasmid, using a 1:1 ratio. Ligated plasmids were transformed to E. cloni® cells (Lucigen) and plated on 28 large agar plates (with antibiotics) in order to conserve library complexity. Approximately ten million colonies were scraped and transferred to an Erlenmeyer for growth.

### OL transcriptional-silencing assay

The oligo-library silencing assay for the transformed OL was developed based on (Sharon et al.) and was carried out as follows:

#### Culture growth

Library-containing bacteria were grown with fresh LB and antibiotic (Kan). Cells were grown to mid-log phase (O.D600 of ~0.6) as measured by a spectrophotometer (Novaspec III, Amersham Biosciences) followed by resuspension with BA buffer and the appropriate antibiotic (Kan). Culture was grown in BA for 3 hours prior to sorting by FACSAria cell sorter (Becton-Dickinson).

#### FACS sorting

Sorting was done at a flow rate of ~20,000 cells per sec. Cells were sorted into 14 bins (500,000 cells per bin) according to the mCherry to eYFP ratio, in two groups: (i) bins 1-8: high resolution on low ratio bins (30% scale), (ii) bins 9-16: full resolution bins (3% scale).

#### Sequencing preparation

Sorted cells were grown overnight in 5 ml LB and appropriate antibiotic (Kan). In the next morning, cells were lysed (TritonX100 0.1% in 1XTE: 15 μl, culture: 5 μl, 99°C for 5 min and 30°C for 5 min) and the DNA from each bin was subjected to PCR with a different 5’ primer containing a specific bin barcode. PCR products were verified in an electrophoresis gel and cleaned using PCR Clean-Up kit (Promega). Equal amounts of DNA (10 ng) from each bin were joined to one 1.5 ml microcentrifuge tube for further analysis.

#### Sequencing

Sample was sequenced on an Illumina Hiseq 2500 Rapid Reagents V2 100 bp paired-end chip. 10% PhiX was added as a control. This resulted in ~140 million reads. NGS processing. From each read, the bin barcode and the sequence of the strain were extracted using a custom Python script consisting of the following steps: paired-end read merge, read orientation fix, identification of the constant parts in the read and extracting the variables: bin barcode, sequence barcode and the variable tested sequence. Finally, each read was mapped to the appropriate combinations of tested sequence and expression bin. This resulted in ~38 million uniquely mapped reads, each containing a perfect match variance sequence and expression bin barcode pair.

#### Inference of per-variant expression profile

We first removed all reads mapped to bin number 16 from the analysis to eliminate biases originating from out-of-range fluorescence measurements. Next, we filtered out sequences with low read counts, keeping only those with a total of at least 10 reads across the bins. This left us with a total of ~36 million reads distributed over 10438 variants. We then generated a single profile by replacing bin 9 with bins 1-8, and redistributing the reads in bin 9 over bins 1-8 according to their relative bin widths. Next, for each sequence we calculated the fraction of cells in each bin, based on the number of sequence reads from that bin that mapped to that variant (the reads of each bin were first normalized to match the fraction of the bin in the entire population). This procedure resulted in expression profiles over 14 bins for 10432 variants (See Supplementary Table 4). The complete Python pipeline is available on Github.

#### Inference of per-variant mean expression level

For each variant we defined the mean expression ratio as the weighted average of the ratios at the geometric centers of the bin, where the weight of each bin is the fraction of the reads from that variant in that bin.

## Position-dependent E5mer effect

To test the effect of E5mer position on the mean e.l.r we fitted the following linear model on the sequences included in Figure 4B:

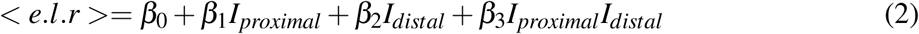

where *I_proximal_* and *I_distal_* are indicators for the presence of proximal (position≥-25 relative to the sigma54 TSS) and distal (position<-25 relative to the sigma54 TSS) E5mers respectively. We then performed a Two-way ANOVA test on the fitted model. The fitted model and the ANOVA tables are presented in Tables S4 and S5.

### Model for translation level with a partially sequestered RBS

#### Classical rate equation model

In the following, we model the bi-phasic nature of mRNA. We begin by considering the simplest model for gene expression. In this model, we assume constant rates for the transcription of RNA *k_r_*, degradation of RNA *γ_r_*, translation of protein from RNA *k_p_*, and the degradation of protein *γ_p_*. This model leads to the following model for mean concentrations of RNA [*R*] and protein [*P*]:

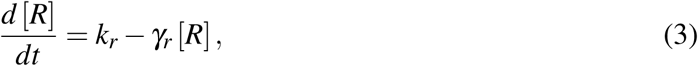

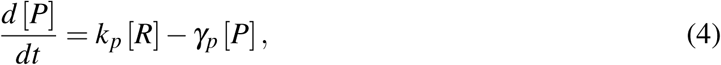

which in steady state leads to the following expression for the mean expression level:

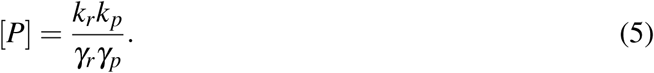

This expression has led to important insights regarding the effect of regulation on gene expression, by showing that regulation can affect the creation constants *k_r_* and *k_p_*, which in turn may result in either an increase or decrease in gene expression (Bintu et al., 2005).

## Adapting the rate equation model to our experiment - the degradation hypothesis

In the case of our experiment, we found that RNA is likely to be found in one of two states or phases: the first is a translationally active “pearled” structure with a low degradation rate, the second is a translationally inactive “branched” structure with a sequestered RBS site and a high degradation rate. Given these observations, the rates as defined above become discrete two-state functions as follows:

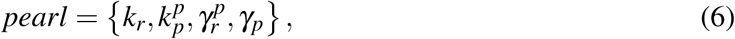

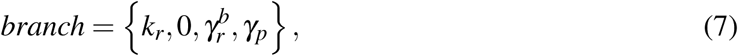

where 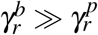.

In order to properly model the resultant gene expression of a bi-phasic state of RNA, we need to first account for the probability of finding a particular RNA molecule in one phase or the other. This probability is sequence dependent, and can be computed using RNA secondary structure computation tools such as Vienna. In general terms, one can think of an ensemble of identical RNA molecules as a collection of secondary structures, each characterized by some energy *ε_i_*. The partition function for this ensemble is:

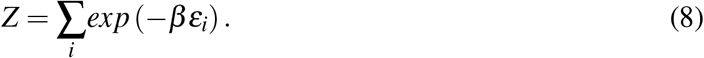

If we now assume that only a certain subset of these configurations exhibits sequestered RBS binding sites, we can express the partition function as a sum of two separate configurational classes:

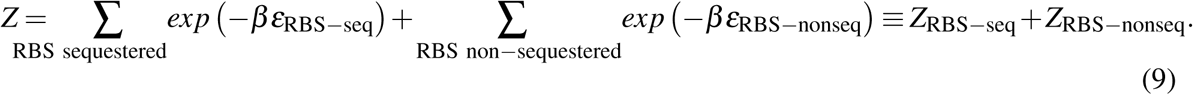

Given this definition of the RNA molecule conifguration space, we can write the following:

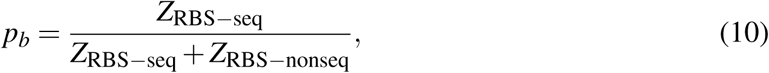

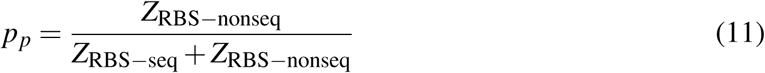

where *p_b_* corresponds to the probability to find the mRNA molecule in a branched phase with a sequestered RBS and *p_p_* represents the probability to find the mRNA molecule in pearled form with multiple ribosomes assembled due to the availability of the RBS.

Given these definitions, we can now rewrite Eqs. 3 and 4 as follows:

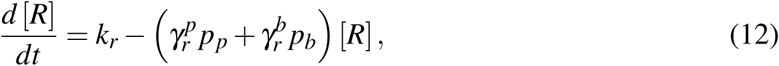

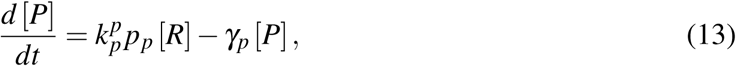

which leads to the following steady state expression for [*P*]:

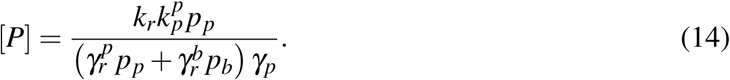

The rate parameters, taken from the BioNumbers database (Milo and Phillips, 2016), were set to 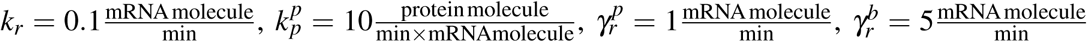 and 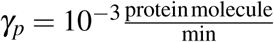.

## Computation of probability for RBS to be sequestered

For each variant, we used the RNAfold tool to calculate the probability of being in the pearled (*p_p_*) and branched (*p_b_* = 1 − *p_p_*) states. According to our hypothesis, an RNA molecule will occupy the pearl phase if the RBS sequence is available for the ribosomes to bind. We defined an available RBS as an RBS sequence in which at least six of the seven bases are unpaired. We th\n calculated the proportion of molecules with available RBS as described in the RNAfold tutorial (https://www.tbi.univie.ac.at/RNA/tutorial/#sec3_2). Briefly, we computed the ensemble free energy of all the chains (β*F_u_* = *−log(Z_RNA_*)),where *Z_R_NA* is the partition function for all the RNA molecule), and assume that all RBS constrained chain are characterized by a typical aSD-SD binding energy (*E_c_*). We then evaluated the probability of molecules with a sequestered RBS using:

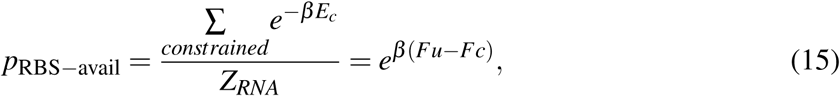

where *F_c_* is the free energy associated with the constrained configurations. Finally, we assumed *p*_RBS−avail_ = *p_p_*.

## Computation of prevalence of CT-rich motifs in sigma54 promoters in *E.coli*

Analysis of the occurrences of aSD:SD around *σ*^54^ TSS positions was carried out in three steps: 1. we first scanned the 50bp region downstream to TSS and locate the first purine-rich hexamer or putative-SD. 2. Next, we searched for the best matching complementary sequence according to a minimal hamming distance (i.e. putative aSD or CT-rich hexmaer) to the purine-rich sequence found in step 1, within 50 bp upstream of the purine-rich hexamer. 3. Finally, tallied the different hamming distance scores to generate the Venn diagram shown in Fig. 6A.

## Algorithm for the assessment of proximal aSD:SD pairs for a bacterial genome

In order to quantitatively assess the prevalence of proximal aSD:SD pairs, our algorithm included the following elements:

### Computation of putative SD hexamers

We constructed a list of all hexamers (4096) and for each one of them we calculated the free energy value (ΔG) for its hybridization with the 16S rRNA of *E.coli* (as in (Li et al., 2012)). We ranked this list of hexamers from low to high free energy value, which corresponds to higher to lower probability for hybridization with the 16S rRNA. The top 20 hexamers were defined as the best 20 putative SD hexamers. (See Table S7).

### Computation of putative aSD hexamers

We computed a set of 20 pyrimidine-rich hexamers with the highest PSSM scores according to our aSD E5mer motif (see Fig. 4A top right for logo). We sorted the list of all hexamers according to their aSD E5mer motif from best aSD (with the highest PSSM value) to least aSD (with the lowest PSSM value). The top 20 hexamers were defined as best 20 aSD hexamers. (See Table S7).

### Computation of random(R) hexamers

We computed a set of 20 randomized hexamers, which did not score highly for either motif. The hexamers were picked at random from the list of 4096. Random hexamers that were found in the top 20 or bottom 20 of the aSD and SD ranked lists were excluded, and replaced by another selection. The same 20 hexamers picked at random for E. coli, were later used for all the other bacterial genomes examined. These are called the random hexamers, denoted R. (See Table S7).

### Computation of proximal aSD:SD and R:SD distributions

Given two sets of 20 hexamers each, A and B, and given one hexamer from each: a∊A and b∊B, we scan the bacterial genome to assess the proximal occurrences of b downstream from a. For every occurrence of a in the genome we asked if we also see b in the d base-pairs downstream from said occurrence of a. The set A can consist of the aSD hexamers or of the R hexamers. The set B consists of the SD hexamers. For each pair of sets, A (aSD or R) and B (always SD), we get 400 pairs of individual elements. For each such pair a:b, we calculated the percentage of proximal pairs (10<d<300) out of all a:b occurrences within 10<d<10000. We thus get a distribution of 400 percentage numbers for the two combination, aSD:SD and R:SD. The scheme is depicted in Fig. S6.

Pseudo-code for the above process:

~~~
Proximal_percentage (G, a, b)
\* a in A=R or aSD, b in SD
ct_prox=0;
ct_total=0;
Find all occurrences of b in G
For each occurrence position p do:
        If a occur in the genomic segment [p−10k, p]
                Then ct_total++;
        If a occur in the genomic segment [p−300, p]
                Then ct_prox++;
return ct_prox/ ct_total
~~~

## Cumulative analysis of aSD:SD depletion in 591 psychrophile and mesophile genomes

We scanned the genomes of additional 591 bacteria (see Supplementary Data 2) For each bacterial genome we compared the percentages of proximal aSD: SD occurrences to the percentages of proximal R:SD occurrences. Each of the bacterial genomes is represented in the scatter plot by the values of x= mean (R:SD), y=mean (aSD:SD).

**Figure S1:**
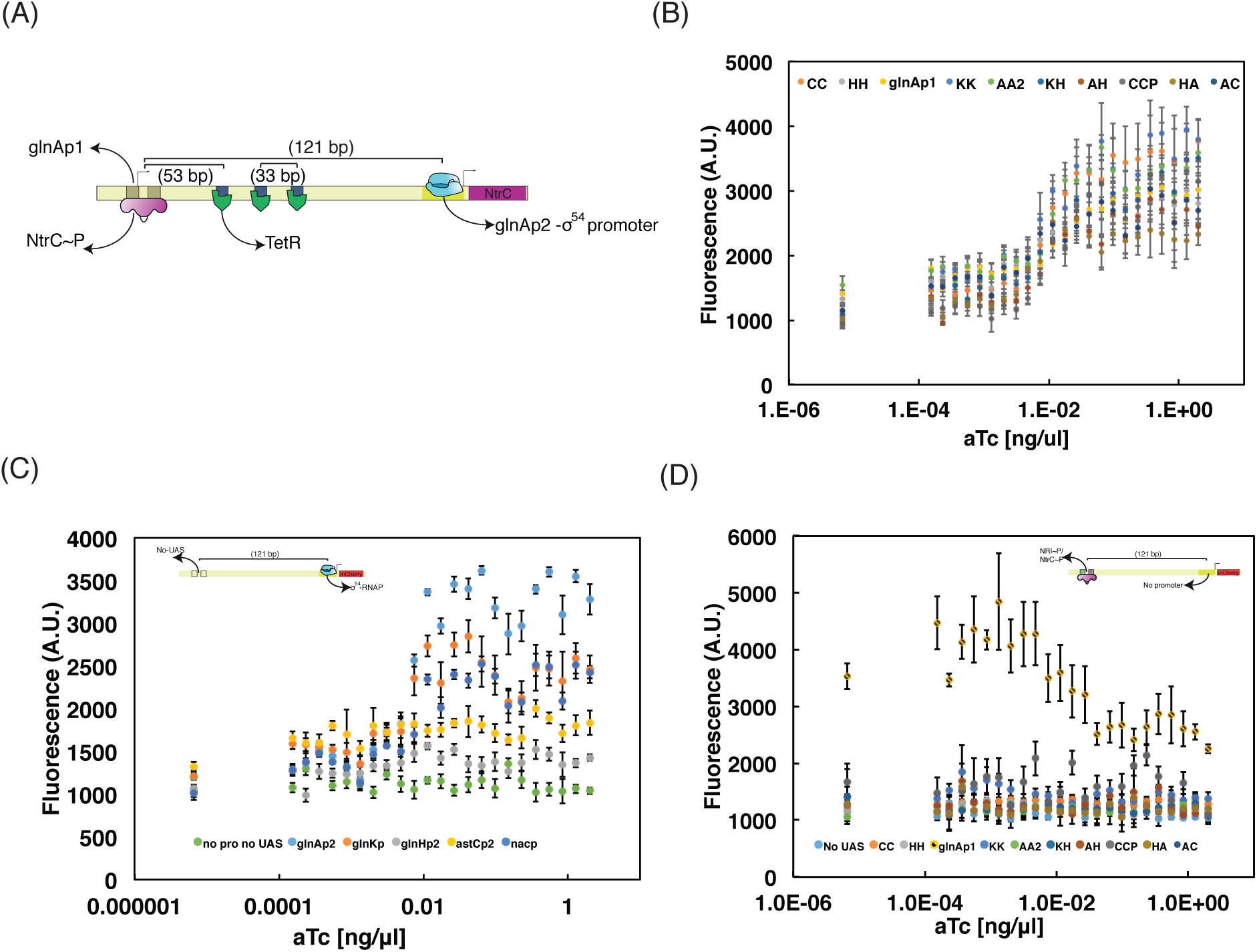
Related to Figure 1. The NtrC switch, cross talk, and control experiments. (A) The NtrC switch circuit design. (B) Sample dose response for one of the promoters (glnKp) with all UASs used in the synthetic enhancer experiment.(C) It was shown that UASs can activate transcription from long distances (Magasanik, 1989). In the experimental design, both the NtrC switch and the UAS-promoter-reporter measurement module were based on *σ*^54^ architecture (thus both containing a UAS and *σ*^54^ promoter) and were located on the same plasmid. We therefore tested whether the NtrC switch’s UAS activates the *σ*^54^ promoter in the measurement module via DNA looping. To achieve this goal, we removed the UAS from the measurement module and measured mCherry fluorescence as a function of aTc concentration. 3 out of 5 promoters (glnAp2, glnKp and nacp) display increasing mCherry fluorescence as a function of increasing aTc concentration, as opposed to glnHp2 and astCp2 that exhibit the same mCherry levels as the no-promoter measurement module (A - light blue). This indicates that cross-activation within our system is possible, depending on the promoter. Interestingly, the two promoters which did not exhibit this effect (glnHp2 and astCp2) are known to be weaker *σ*^54^ promoters (Reitzer and Schneider, 2001). (D) In order to verify that the mCherry expression observed in the experiment originated from the tested *σ*^54^ promoter in the measurement module, we removed the promoter from the measurement module and measured mCherry fluorescence. As shown in the panel, the glnAp1 UAS, which contains a *σ*^70^ promoter, was the only UAS for which mCherry was expressed. Fluorescence results for glnAp1 show up to two-fold repression at maximal aTc concentration, consistent with the fact that glnAp1 acts as not only as a UAS but also as a promoter. As NtrC concentrations increase, more NtrC binds to the glnAp1 UAS and the glnAp1 promoter activity is repressed (Reitzer and Magasanik 1985).

**Figure S2:**
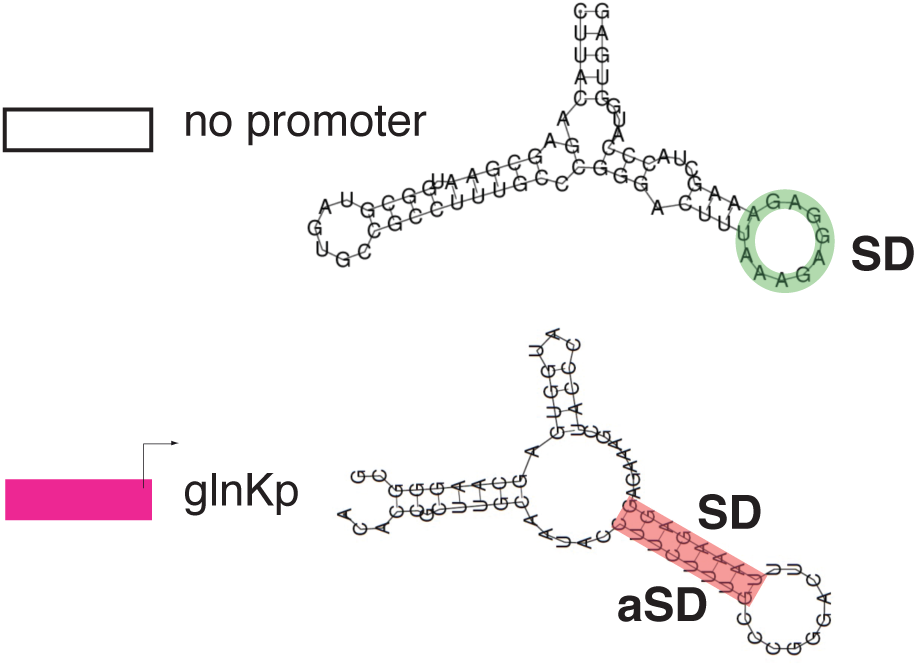
Related to Figure 2. Putative insulator secondary structure. RNAfold generated secondary structures for the no σ54-dependent promoter control (top: -46 to +24 – with 0 defined as the TSS), and the glnKp construct (bottom: -38 to +32 with 0 definedas the TSS) highlighting in green the RBS or Shine Dalgarno (SD) sequences for both structures. In the two case, SD sequence is either single stranded (Top), or sequestered in a hairpin structure (Bottom).

**Figure S3:**
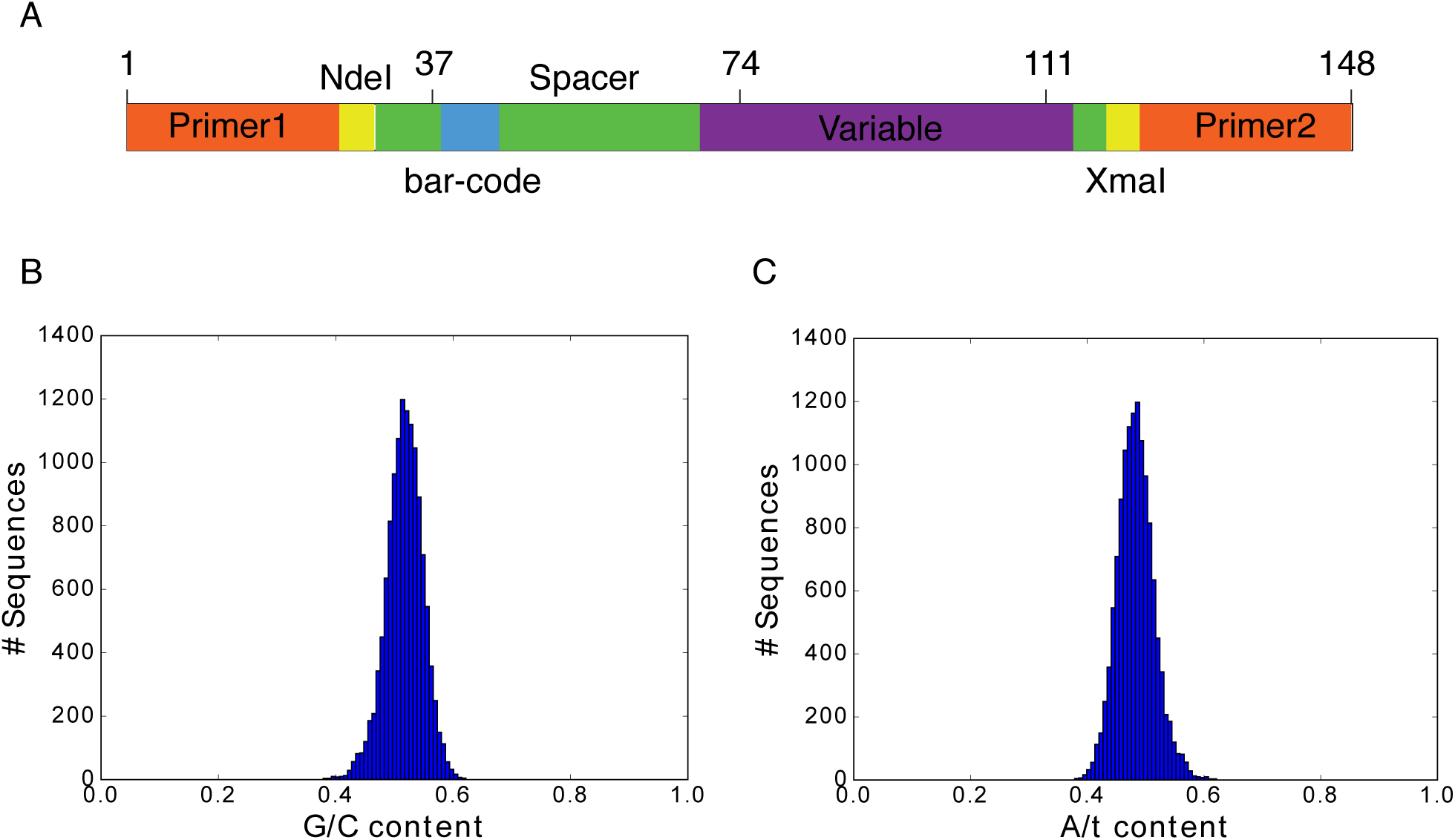
Related to Figure 3. OL structure. (A) The library contained 12758 unique sequences, each of length 145-148 bp, with the following structure: red - universal primers for amplification (44 bp), orange- restriction sites (12 bp), green - universal spacers (32 bp), blue #1 - variable barcode (10 bp), blue #2 - variable (*σ*^54^ promoter) region (47-50 bp). (B) GC content distribution. (C) AT content distribution.

**Figure S4:**
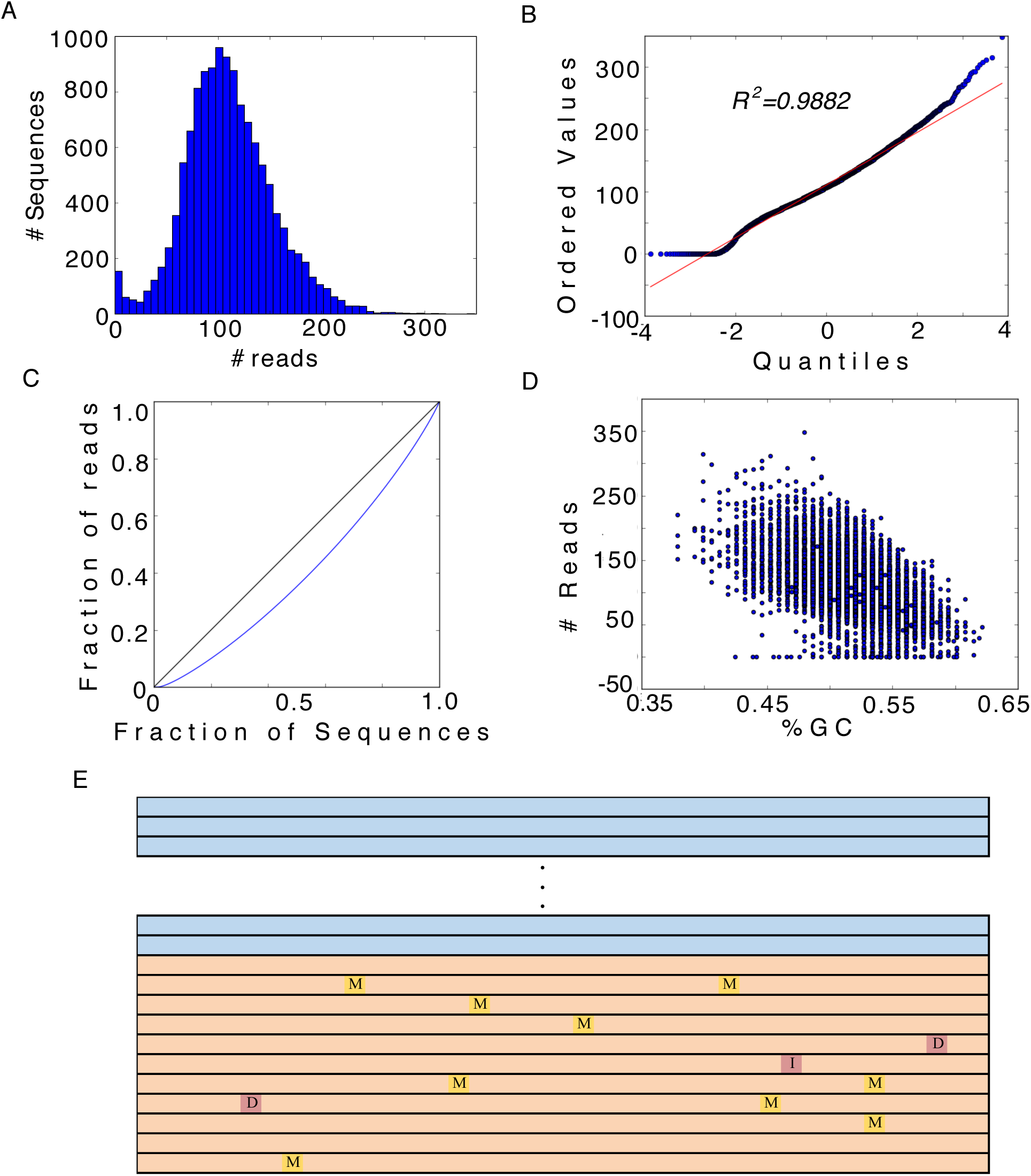
Related to Figure 3. Variant abundance analysis. (A) Read count histogram showing a mean of 110 reads per variant and a SD of 43. (B) Probability plot against the normal distribution. (C) Lorentz curve. (D) Effect of GC content on variant abundance. (E) Schematic of the position of errors along variant length.

**Figure S5:**
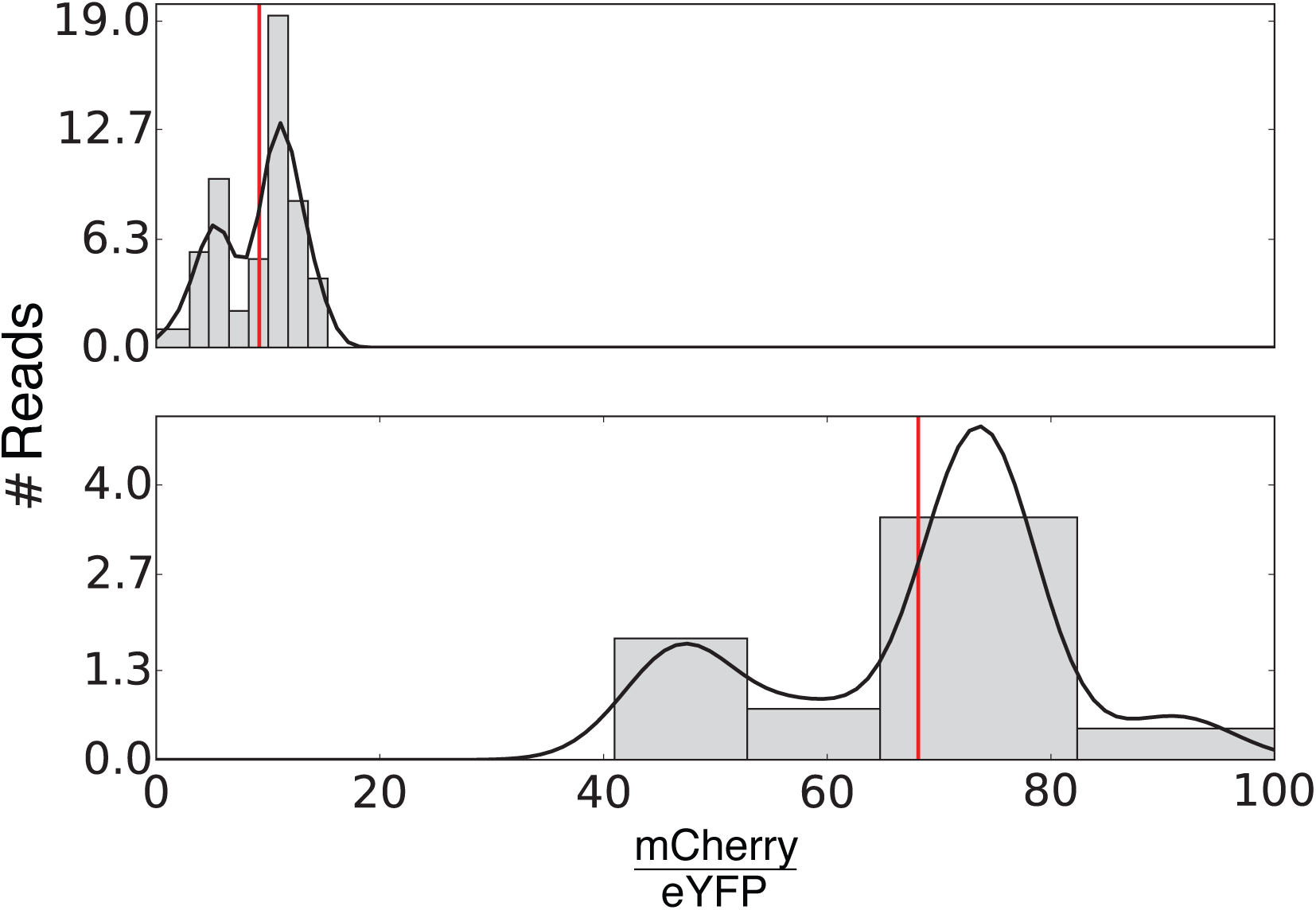
Related to Figure 3. Single sequence variant expression profile: sample data showing the number of reads as a function of mean fluorescence ratio obtained for silencing (top) and non-silencing (bottom) variants, respectively. Straight lines correspond to a smoothing procedure done with a cubic-spline fit to the data.

**Figure S6:**
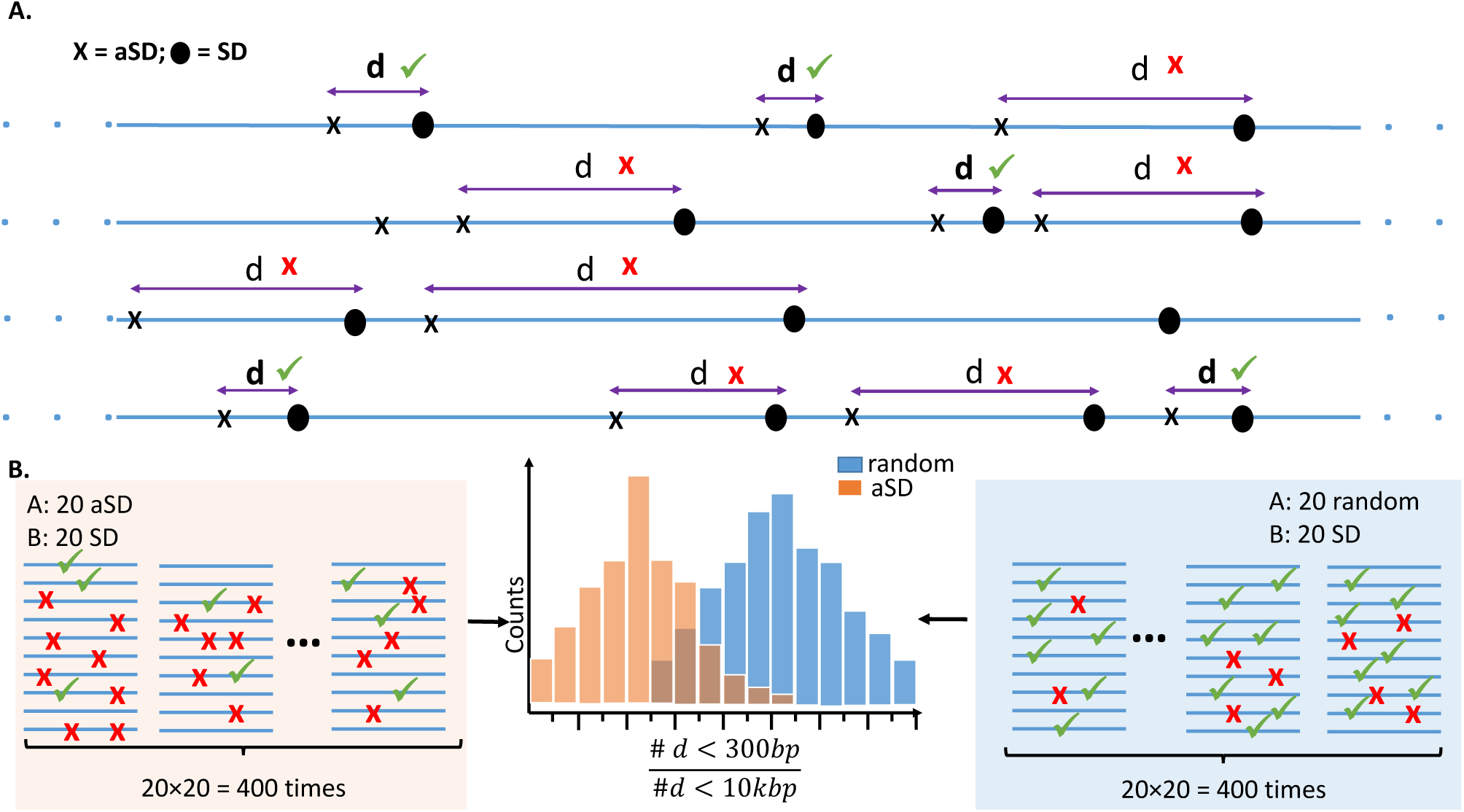
Related to Figure 5. Scheme of aSD:SD vs. random:SD proximal occurrences analysis. A. Distances between SD and upstream aSD are measured. The percentage of proximal occurrences within less than 300 bp (marked with green check mark) is calculated with respect to all aSD:SD occurrences within 10kbp (both red Xs and green check marks). B. 20×20=400 aSD:SD occurrence percentages (left, marked with orange shaded box) and 400 R:SD (right, marked with blue shaded box) occurrence percentages are compared. P-values are calculated using Wilcoxon test.

**Table S1:**
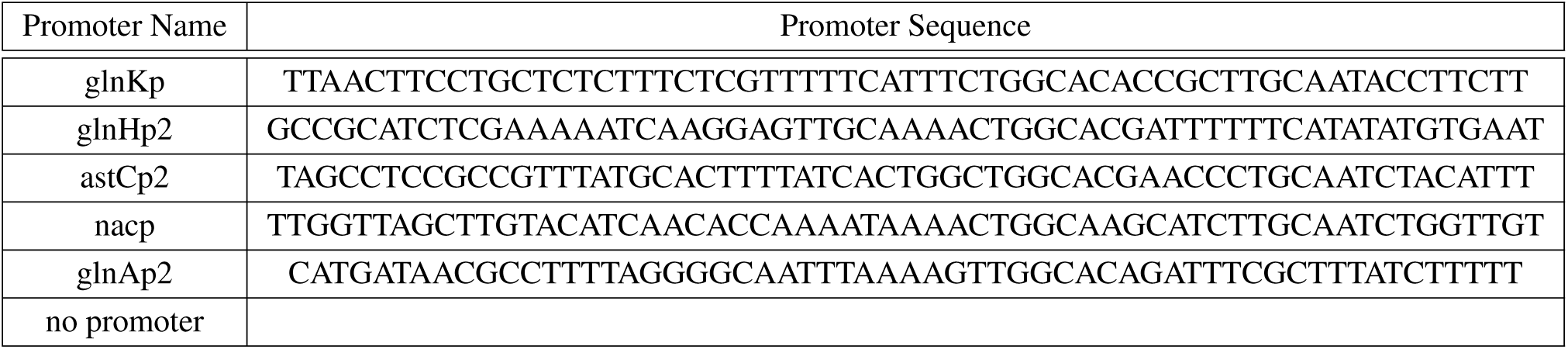
*σ*^54^ promoters used in the UAS-promoter-reporter experiment. Related to Figure 1.

**Table S2:**
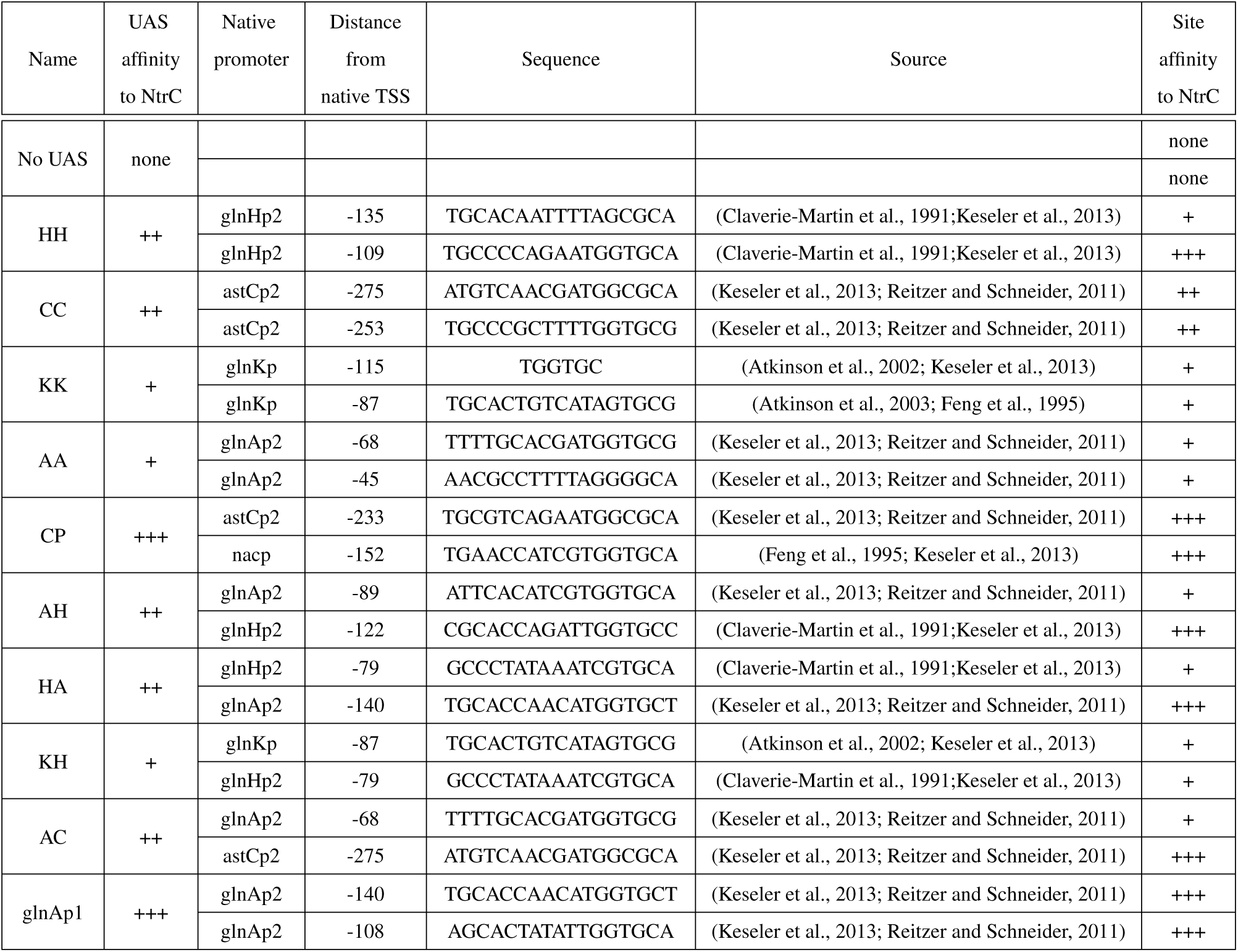
Synthetic and natural UAS sequences used in the NtrC switch. Related to Figure 1.

**Table S3:**
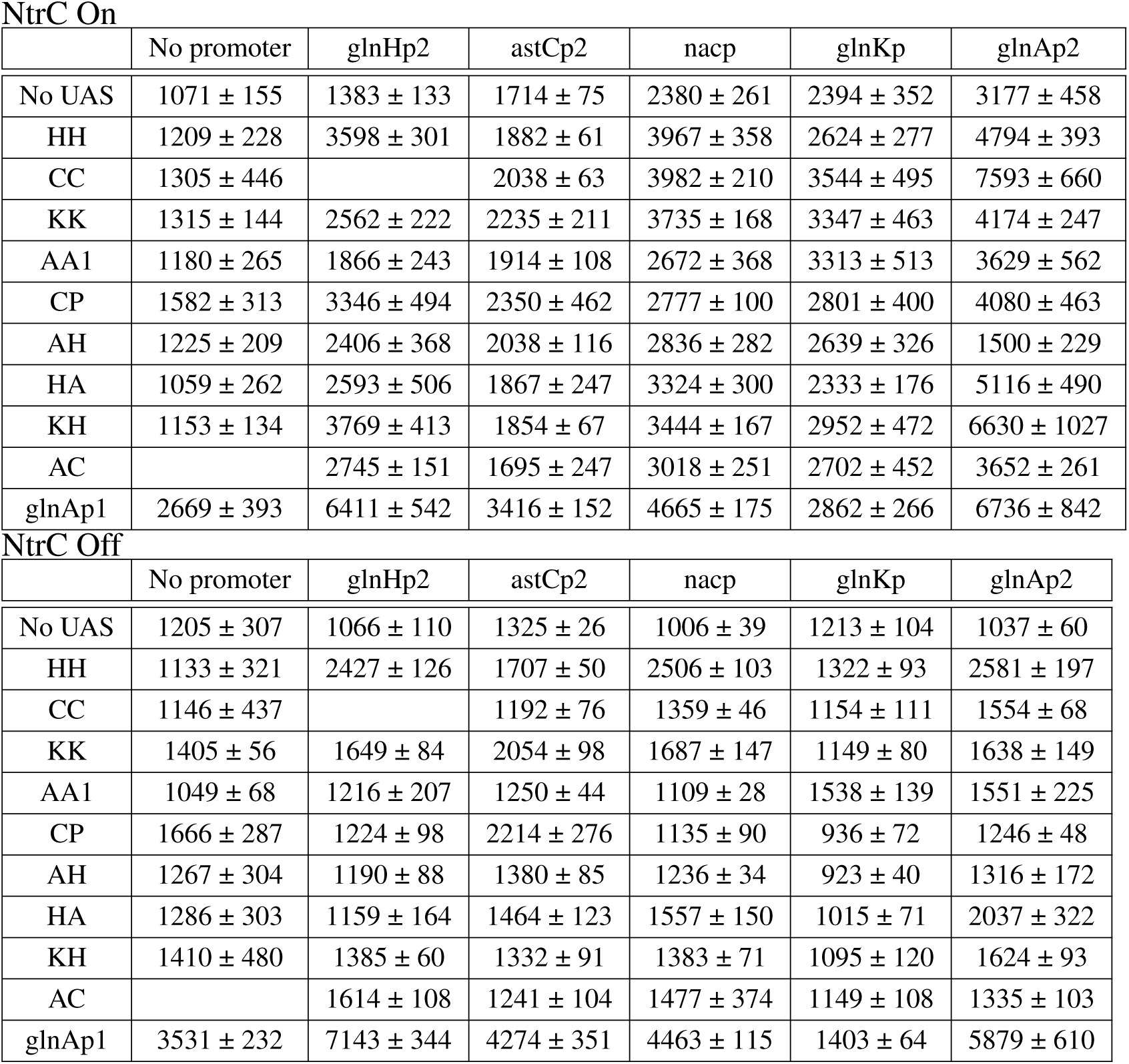
Mean fluorescence expression-level data in steady state for the synthetic enhancers together with their variation. Related to Fig. 1B.

NtrC On
|  | No promoter | glnHp2 | astCp2 | nacp | glnKp | glnAp2 |
| --- | --- | --- | --- | --- | --- | --- |
| No UAS | 1071 ± 155 | 1383 ± 133 | 1714 ± 75 | 2380 ± 261 | 2394 ± 352 | 3177 ± 458 |
| HH | 1209 ± 228 | 3598 ± 301 | 1882 ± 61 | 3967 ± 358 | 2624 ± 277 | 4794 ± 393 |
| CC | 1305 ± 446 |  | 2038 ± 63 | 3982 ± 210 | 3544 ± 495 | 7593 ± 660 |
| KK | 1315 ± 144 | 2562 ± 222 | 2235 ± 211 | 3735 ± 168 | 3347 ± 463 | 4174 ± 247 |
| AA1 | 1180 ± 265 | 1866 ± 243 | 1914 ± 108 | 2672 ± 368 | 3313 ± 513 | 3629 ± 562 |
| CP | 1582 ± 313 | 3346 ± 494 | 2350 ± 462 | 2777 ± 100 | 2801 ± 400 | 4080 ± 463 |
| AH | 1225 ± 209 | 2406 ± 368 | 2038 ± 116 | 2836 ± 282 | 2639 ± 326 | 1500 ± 229 |
| HA | 1059 ± 262 | 2593 ± 506 | 1867 ± 247 | 3324 ± 300 | 2333 ± 176 | 5116 ± 490 |
| KH | 1153 ± 134 | 3769 ± 413 | 1854 ± 67 | 3444 ± 167 | 2952 ± 472 | 6630 ± 1027 |
| AC |  | 2745 ± 151 | 1695 ± 247 | 3018 ± 251 | 2702 ± 452 | 3652 ± 261 |
| glnAp1 | 2669 ± 393 | 6411 ± 542 | 3416 ± 152 | 4665 ± 175 | 2862 ± 266 | 6736 ± 842 |

|  | No promoter | glnHp2 | astCp2 | nacp | glnKp | glnAp2 |
| --- | --- | --- | --- | --- | --- | --- |
| No UAS | 1205 ± 307 | 1066 ± 110 | 1325 ± 26 | 1006 ± 39 | 1213 ± 104 | 1037 ± 60 |
| HH | 1133 ± 321 | 2427 ± 126 | 1707 ± 50 | 2506 ± 103 | 1322 ± 93 | 2581 ± 197 |
| CC | 1146 ± 437 |  | 1192 ± 76 | 1359 ± 46 | 1154 ± 111 | 1554 ± 68 |
| KK | 1405 ± 56 | 1649 ± 84 | 2054 ± 98 | 1687 ± 147 | 1149 ± 80 | 1638 ± 149 |
| AA1 | 1049 ± 68 | 1216 ± 207 | 1250 ± 44 | 1109 ± 28 | 1538 ± 139 | 1551 ± 225 |
| CP | 1666 ± 287 | 1224 ± 98 | 2214 ± 276 | 1135 ± 90 | 936 ± 72 | 1246 ± 48 |
| AH | 1267 ± 304 | 1190 ± 88 | 1380 ± 85 | 1236 ± 34 | 923 ± 40 | 1316 ± 172 |
| HA | 1286 ± 303 | 1159 ± 164 | 1464 ± 123 | 1557 ± 150 | 1015 ± 71 | 2037 ± 322 |
| KH | 1410 ± 480 | 1385 ± 60 | 1332 ± 91 | 1383 ± 71 | 1095 ± 120 | 1624 ± 93 |
| AC |  | 1614 ± 108 | 1241 ± 104 | 1477 ± 374 | 1149 ± 108 | 1335 ± 103 |
| glnAp1 | 3531 ± 232 | 7143 ± 344 | 4274 ± 351 | 4463 ± 115 | 1403 ± 64 | 5879 ± 610 |

**Table S4:** Synthesis error rates by type and base. Related to Figure 3.

|  | Mismatch rate<br>[1/bases] | Deletion (1 nt)<br>[1/bases] | Insertion (1 nt)<br>[rate 1/bases] | Deletion (>1 nt)<br>[1/reads] | Insertion (>1 nt)<br>[1/reads] |
| --- | --- | --- | --- | --- | --- |
| A | 2713 | 1682 | 5461 | 1138 | 5365 |
| C | 2135 | 1614 | 4310 | 1205 | 7950 |
| T | 1946 | 2036 | 5777 | 1322 | 6456 |
| G | 789 | 1469 | 5367 | 1153 | 6287 |
| Average | 1550 | 1671 | 5133 | 1198 | 6402 |

**Table S5:**
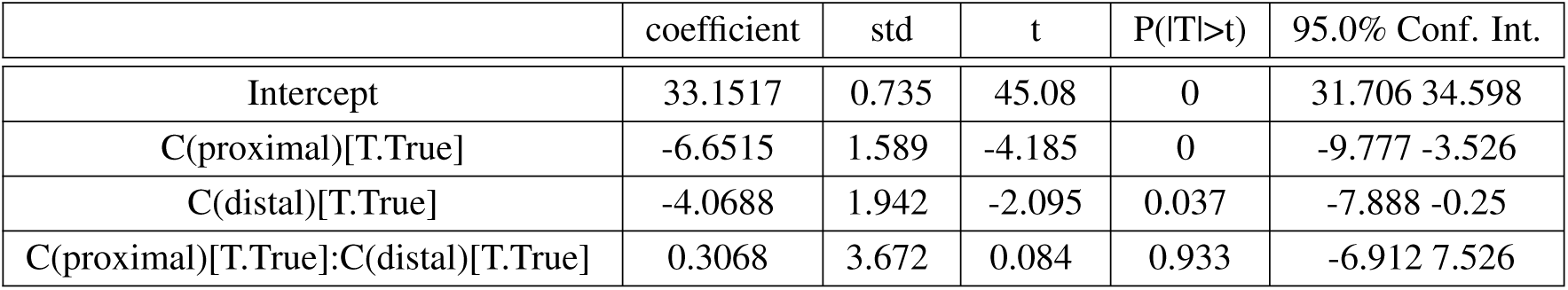
Summary of the fitted model for the additivity of the insulating effects of proximal and distal CT-rich motifs. Related to Figure 4.

**Table S6:** A two-way ANOVA test on the fitted model. Related to Figure 4.

|  | sum_sq | df | f | P(F>f) |
| --- | --- | --- | --- | --- |
| C(proximal) | 2943.800 | 1 | 21.180 | 0.000006 |
| C(distal) | 811.527 | 1 | 5.839 | 0.016 |
| C(proximal):C(distal) | 0.970 | 1 | 0.007 | 0.93 |
| Residual | 53372.028 | 384 |  |  |

**Table S7:** Ranked list of SD, aSD, and Random hexamer. Related to Figure 6.

| The best SD |  |  | The best aSD |  |  | Random |  |  |
| --- | --- | --- | --- | --- | --- | --- | --- | --- |
| Seq | $\Delta G$ | rank | Seq | E5mer_PSSM | rank | Seq | E5mer_pssm | rank |
| GGAGGT | -10.2 | 1 | TCCCCC | -1.08992 | 1 | CTCTTG | -3.20252 | 617 |
| GGAGGG | -9.8 | 2 | CCCCCC | -1.16622 | 2 | TCATTT | -3.24237 | 665 |
| GGAGGA | -9.3 | 3 | TCCCCT | -1.18705 | 3 | CCACAC | -3.32221 | 765 |
| GGAGGC | -9.3 | 4 | CCCCCT | -1.26335 | 4 | CACACC | -3.33037 | 774 |
| GGGGGT | -9.3 | 5 | TCCCTC | -1.36079 | 5 | CTCACA | -3.36743 | 814 |
| AGGAGG | -9.1 | 6 | CCCCTC | -1.4371 | 6 | GTCTCG | -3.37205 | 819 |
| GGGAGG | -9.1 | 7 | TCCCTT | -1.45792 | 7 | CCCAAC | -3.4567 | 929 |
| CGGAGG | -8.9 | 8 | CCCCTT | -1.53423 | 8 | ATCTCA | -3.48297 | 962 |
| GGGGGG | -8.9 | 9 | TTCCCC | -1.57058 | 9 | ACCGTG | -3.51886 | 1006 |
| GAGGTG | -8.8 | 10 | TCGCCC | -1.58113 | 10 | GACCAC | -3.70653 | 1210 |
| TGGAGG | -8.7 | 11 | TGCCCC | -1.60422 | 11 | CTCGGT | -3.84439 | 1437 |
| GGGGGA | -8.4 | 12 | GCCCCC | -1.60663 | 12 | TGTTCA | -4.01378 | 1740 |
| GGGGGC | -8.4 | 13 | TCCCCG | -1.6331 | 13 | TAGCGA | -4.22422 | 2015 |
| AGGGGG | -8.2 | 14 | TCCCCA | -1.6331 | 14 | TTACGA | -4.26976 | 2097 |
| CGGGGG | -8.0 | 15 | CTCCCC | -1.64689 | 15 | GAGCGT | -4.29488 | 2139 |
| GGGGTG | -7.9 | 16 | CCGCCC | -1.65744 | 16 | CTTACG | -4.49223 | 2508 |
| TGGGGG | -7.8 | 17 | TTCCCT | -1.66771 | 17 | CGATGC | -4.5781 | 2655 |
| GAGGTA | -7.0 | 18 | TCGCCT | -1.67826 | 18 | AGTACT | -4.63116 | 2726 |
| AGGAGA | -6.8 | 19 | CGCCCC | -1.68053 | 19 | TTGAAA | -4.89546 | 3067 |
| CGAGGT | -6.8 | 20 | TGCCCT | -1.70136 | 20 | ATTTAG | -5.7209 | 3891 |

## References

Amit, R., Garcia, H.G., Phillips, R., and Fraser, S.E. (2011). Building Enhancers from the Ground Up: A Synthetic Biology Approach. Cell 146, 105–118.

Atkinson, M.R., Blauwkamp, T.A., Bondarenko, V., Studitsky, V., and Ninfa, A.J. (2002). Activation of the glnA, glnK, and nac Promoters as Escherichia coli Undergoes the Transition from Nitrogen Excess Growth to Nitrogen Starvation. J. Bacteriol. 184, 5358–5363.

Atkinson, M.R., Savageau, M.A., Myers, J.T., and Ninfa, A.J. (2003). Development of Genetic Circuitry Exhibiting Toggle Switch or Oscillatory Behavior in Escherichia coli. Cell 113, 597–607.

Babitzke, P., Baker, C.S., and Romeo, T. (2009). Regulation of Translation Initiation by RNA Binding Proteins. Annu. Rev. Microbiol. 63, 27–44.

Barrios, H., Valderrama, B., and Morett, E. (1999). Compilation and analysis of σ54-dependent promoter sequences. Nucleic Acids Res. 27, 4305–4313.

Bentele, K., Saffert, P., Rauscher, R., Ignatova, Z., and Blüthgen, N. (2013). Efficient translation initiation dictates codon usage at gene start. Mol. Syst. Biol. 9, 675.

Bintu, L., Buchler, N. E., Garcia, H. G., Gerland, U., Hwa, T., Kondev, J., Kuhlman, T. and Phillips, R. (2005). Transcriptional regulation by the numbers: applications. Current Opinion in Genetics & Development 15, 125–135.

Bonocora, R.P., Smith, C., Lapierre, P., and Wade, J.T. (2015). Genome-Scale Mapping of Escherichia coli σ54 Reveals Widespread, Conserved Intragenic Binding. PLoS Genet. 11.

Brunwasser-Meirom, M., Pollak, Y., Goldberg, S., Levy, L., Atar, O. and Amit, R. (2016). Using synthetic bacterial enhancers to reveal a looping-based mechanism for quenching-like repression. Nature Communications 7, 10407.

Calin-Jageman, I., and Nicholson, A.W. (2003). Mutational Analysis of an RNA Internal Loop as a Reactivity Epitope for Escherichia coli Ribonuclease III Substrates. Biochemistry (Mosc.) 42, 5025–5034.

Campo, C.D., Bartholomäus, A., Fedyunin, I., and Ignatova, Z. (2015). Secondary Structure across the Bacterial Transcriptome Reveals Versatile Roles in mRNA Regulation and Function. PLOS Genet 11, e1005613.

Claverie-Martin, F., and Magasanik, B. (1991). Role of integration host factor in the regulation of the glnHp2 promoter of Escherichia coli. Proc. Natl. Acad. Sci. 88, 1631–1635.

Deana, A., Celesnik, H., and Belasco, J.G. (2008). The bacterial enzyme RppH triggers messenger RNA degradation by 5′ pyrophosphate removal. Nature 451, 355–358.

De Smit, M.H., and Duin, J.V. (1990). Control of Prokaryotic Translational Initiation by mRNA Secondary Structure. In Progress in Nucleic Acid Research and Molecular Biology, W.E.C. and K. Moldave, ed. (Academic Press), pp. 1–35.

Eden, E., Lipson, D., Yogev, S., and Yakhini, Z. (2007). Discovering Motifs in Ranked Lists of DNA Sequences. PLOS Comput Biol 3, e39.

Epshtein, V., Toulmé, F., Rahmouni, A.R., Borukhov, S., and Nudler, E. (2003). Transcription through the roadblocks: the role of RNA polymerase cooperation. EMBO J. 22, 4719–4727.

Farley, E.K., Olson, K.M., Zhang, W., Brandt, A.J., Rokhsar, D.S., and Levine, M.S. (2015). Suboptimization of developmental enhancers. Science 350, 325–328.

Feng, J., Goss, T.J., Bender, R.A., and Ninfa, A.J. (1995). Repression of the Klebsiella aerogenes nac promoter. J. Bacteriol. 177, 5535–5538.

Francke, C., Groot Kormelink, T., Hagemeijer, Y., Overmars, L., Sluijter, V., Moezelaar, R., and Siezen, R.J. (2011). Comparative analyses imply that the enigmatic sigma factor 54 is a central controller of the bacterial exterior. BMC Genomics 12, 385.

Gu, W., Zhou, T., and Wilke, C.O. (2010). A Universal Trend of Reduced mRNA Stability near the Translation-Initiation Site in Prokaryotes and Eukaryotes. PLOS Comput Biol 6, e1000664.

Hao, N., Palmer, A.C., Ahlgren-Berg, A., Shearwin, K.E., and Dodd, I.B. (2016). The role of repressor kinetics in relief of transcriptional interference between convergent promoters. Nucleic Acids Res. 44, 6625–6638.

Hofacker, I.L., Fontana, W., Stadler, P.F., Bonhoeffer, L.S., Tacker, M., and Schuster, P. (1994). Fast folding and comparison of RNA secondary structures. Monatshefte Für Chem. Chem. Mon. 125, 167–188.

Hoover, T.R., Santero, E., Porter, S., and Kustu, S. (1990). The integration host factor stimulates interaction of RNA polymerase with NIFA, the transcriptional activator for nitrogen fixation operons. Cell 63, 11–22.

Hui, M.P., Foley, P.L., and Belasco, J.G. (2014). Messenger RNA Degradation in Bacterial Cells. Annu. Rev. Genet. 48, 537–559.

Jakobsen, J.S., Jelsbak, L., Jelsbak, L., Welch, R.D., Cummings, C., Goldman, B., Stark, E., Slater, S., and Kaiser, D. (2004). σ54 Enhancer Binding Proteins and Myxococcus xanthus Fruiting Body Development. J. Bacteriol. 186, 4361–4368.

J. Schaefer, C. Engl, N. Zhang, E. Lawton, and M. Buck (2015). Genome wide interactions of wild-type and activator bypass forms of 54. Nucleic Acids Res.

Keseler, I. M., Mackie, A., Peralta-Gil, M., Santos-Zavaleta, A., Gama-Castro, S., Bonavides-Martínez, C., Fulcher, C., Huerta, A. M., Kothari, A., Krummenacker, M., Latendresse, M., Muñiz Rascado, L., Ong, Q., Paley, S., Schröder, I., Shearer, A. G., Subhraveti, P., Travers, M., Weerasinghe, D., Weiss, V., Collado-Vides, J., Gunsalus, R. P., Paulsen, I. and Karp, P. D. (2013). EcoCyc: fusing model organism databases with systems biology. Nucleic Acids Research 41, D605-D612.

Kinney, J.B., Murugan, A., Callan, C.G., and Cox, E.C. (2010). Using deep sequencing to characterize the biophysical mechanism of a transcriptional regulatory sequence. Proc. Natl. Acad. Sci. U. S. A. 107, 9158–9163.

Kiupakis, A.K., and Reitzer, L. (2002). ArgR-Independent Induction and ArgR-Dependent Superinduction of the astCADBE Operon in Escherichia coli. J. Bacteriol. 184, 2940–2950.

Komarova, A.V., Tchufistova, L.S., Supina, E.V., and Boni, I.V. (2002). Protein S1 counteracts the inhibitory effect of the extended Shine-Dalgarno sequence on translation. RNA 8, 1137–1147.

Korbel, J.O., Jensen, L.J., von Mering, C., and Bork, P. (2004). Analysis of genomic context: prediction of functional associations from conserved bidirectionally transcribed gene pairs. Nat. Biotechnol. 22, 911–917.

Kudla, G., Murray, A.W., Tollervey, D., and Plotkin, J.B. (2009). Coding-sequence determinants of gene expression in Escherichia coli. Science 324, 255–258.

Leibovich, L., and Yakhini, Z. (2012). Efficient motif search in ranked lists and applications to variable gap motifs. Nucleic Acids Res. gks206.

Leibovich, L., Paz, I., Yakhini, Z., and Mandel-Gutfreund, Y. (2013). DRIMust: a web server for discovering rank imbalanced motifs using suffix trees. Nucleic Acids Res. 41, W174–W179.

Li, G.-W., Oh, E., and Weissman, J.S. (2012). The anti-Shine-Dalgarno sequence drives translational pausing and codon choice in bacteria. Nature 484, 538–541.

Ma, J., Campbell, A., and Karlin, S. (2002). Correlations between Shine-Dalgarno Sequences and Gene Features Such as Predicted Expression Levels and Operon Structures. J. Bacteriol. 184, 5733–5745.

Mackie, G.A. (1998). Ribonuclease E is a 5′-end-dependent endonuclease. Nature 395, 720–724.

Magasanik, B. (1989). Gene regulation from sites near and far. The New Biologist 1, 247–251.

Milo & Philips, R.& R. (2016). Cell Biology by the Numbers (Garland Science).

Na, D., and Lee, D. (2010). RBSDesigner: software for designing synthetic ribosome binding sites that yields a desired level of protein expression. Bioinformatics 26, 2633–2634.

Nakamoto, T. (2006). A unified view of the initiation of protein synthesis. Biochem. Biophys. Res. Commun. 341, 675–678.

Peterman, N., and Levine, E. (2016). Sort-seq under the hood: implications of design choices on large-scale characterization of sequence-function relations. BMC Genomics 17, 206.

Reitzer, L., and Schneider, B.L. (2001). Metabolic context and possible physiological themes of sigma(54)-dependent genes in Escherichia coli. Microbiol. Mol. Biol. Rev. MMBR 65, 422–444, table of contents.

Reitzer, L. J. and Magasanik, B. (1985). Expression of glnA in Escherichia coli is regulated at tandem promoters. Proceedings of the National Academy of Sciences USA 82, 1979–1983.

Richards, J., Luciano, D.J., and Belasco, J.G. (2012). Influence of translation on RppH-dependent mRNA degradation in Escherichia coli. Mol. Microbiol. 86, 1063–1072.

Robertson, H.D. (1982). Escherichia coli ribonuclease III cleavage sites. Cell 30, 669–672.

Salgado, H., Peralta-Gil, M., Gama-Castro, S., Santos-Zavaleta, A., Muniz-Rascado, L., Garcia-Sotelo, J.S., Weiss, V., Solano-Lira, H., Martinez-Flores, I., Medina-Rivera, A., et al. (2013). RegulonDB v8.0: omics data sets, evolutionary conservation, regulatory phrases, cross-validated gold standards and more. Nucleic Acids Res. 41, D203–D213.

Scharff, L.B., Childs, L., Walther, D., and Bock, R. (2011). Local Absence of Secondary Structure Permits Translation of mRNAs that Lack Ribosome-Binding Sites PLOS Genet. 7, e1002155.

Schwab, D., and Bruinsma, R.F. (2009). Flory Theory of the Folding of Designed RNA Molecules. J. Phys. Chem. B 113, 3880–3893.

Schwartz, M., Roa, M., and Débarbouillé, M. (1981). Mutations that affect lamB gene expression at a posttranscriptional level. Proc. Natl. Acad. Sci. U. S. A. 78, 2937–2941.

Shalem, O., Sharon, E., Lubliner, S., Regev, I., Lotan-Pompan, M., Yakhini, Z., and Segal, E. (2015). Systematic Dissection of the Sequence Determinants of Gene 3’ End Mediated Expression Control. PLOS Genet 11, e1005147.

Sharon, E., Kalma, Y., Sharp, A., Raveh-Sadka, T., Levo, M., Zeevi, D., Keren, L., Yakhini, Z., Weinberger, A., and Segal, E. (2012). Inferring gene regulatory logic from high-throughput measurements of thousands of systematically designed promoters. Nat. Biotechnol. 30, 521–530.

Sharon, E., Dijk, D. van, Kalma, Y., Keren, L., Manor, O., Yakhini, Z., and Segal, E. (2014). Probing the effect of promoters on noise in gene expression using thousands of designed sequences. Genome Res. gr. 168773.113.

Shearwin, K.E., Callen, B.P., and Egan, J.B. (2005). Transcriptional interference - a crash course. Trends Genet. 21, 339–345.

Winkler, W.C., and Breaker, R.R. (2005). Regulation of Bacterial Gene Expression by Riboswitches. Annu. Rev. Microbiol. 59, 487–517.

Yokobayashi, Y., Weiss, R., and Arnold, F.H. (2002). Directed evolution of a genetic circuit. Proc. Natl. Acad. Sci. 99, 16587–16591.

